# Platelet-derived CXCL7 induces neutrophil extracellular traps via CXCL7/CXCR2 axis, exacerbating the pathogenesis of diabetic retinopathy

**DOI:** 10.64898/2026.01.22.701201

**Authors:** Bishamber Nath, Suraj Bhausaheb Mungase, Ritu Sharaya, Anshu Gupta, Amir Ali, Mahesh J Kulkarni, Manabjyoti Barman, Sudhagar Selvaraj, Amit Kumar Yadav, Ramu Adela

**Affiliations:** Department of Pharmacy Practice, National Institute of Pharmaceutical Education and Research (NIPER) Guwahati, Sila Katamur, Halugurisuk, Changsari, Kamrup, Assam, PIN-781101, India; Computational and Mathematical Biology Centre, Translational Health Science and Technology Institute, NCR, Biotech Science Cluster, 3^rd^ Milestone, Faridabad-Gurgaon Expressway, Faridabad, Haryana-121001, India; Department of Biotechnology, National Institute of Pharmaceutical Education and Research (NIPER) Guwahati, Sila Katamur, Halugurisuk, Changsari, Kamrup, Assam, PIN-781101, India; Proteomics Facility, Biochemical Sciences Division, CSIR-National Chemical Laboratory, Pune 411008, India; Deaprtment of Vitreo-retina, Sri Sankaradeva Nethralaya, Guwahati, Assam, 781028, India

**Keywords:** Platelet, Neutrophil, Neutrophil extracellular traps (NETs), CXCL7, CXCR2, Diabetic retinopathy

## Abstract

**Background:** Prolonged hyperglycemia in diabetes activates platelets and immune cells, forming platelet-immune complexes that damage blood vessels in the retina. However, the role of platelet-neutrophil interactions and neutrophil extracellular traps(NETs) in the development of diabetic retinopathy (DR) was not well studied. In this study, we investigated the mechanisms underlying platelet-mediated NET formation in DR.

**Methodology:** Platelet activation markers, platelet-neutrophil aggregates (PNA), and NETs markers were assessed by flowcytometry, and the circulatory level of inflammatory markers was measured by Luminex assays in healthy control(HC), type 2 diabetes mellitus(T2DM), non-proliferative DR(NPDR) and proliferative DR(PDR) subjects. In vitro studies investigated platelet-neutrophil interaction in NETs formation using an immunofluorescence assay. Proteomics analysis identified the mechanistic regulators of platelet-induced NETs in DR. Platelet pellet and plasma CXCL7 were quantified using western blot and ELISA, respectively. The role of the CXCL7/CXCR2 axis in inducing NETs formation was examined using CXCL7 recombinant protein, anti-CXCL7 antibody and CXCR2 antagonist (SB225002).

**Results:** Platelet activation markers (p-selectin & PF4), PNA, and NETs markers (%NETs, proteinase-3 (PR3), neutrophil elastase (NE)) were significantly increased in the DR group. In vitro studies confirmed that DR-platelets aggregate with healthy neutrophils and form NETs compared to T2DM and HC-platelets. Furthermore, platelet activation and NETs markers were positively correlated with pro-angiogenic (ANGPT2, VEGFA) and inflammatory markers (IL18, ICAM1). In vitro studies reveal that NETs induce inflammation, endothelial dysfunction, disrupt the endothelial monolayer and exacerbate angiogenesis in RF/6A endothelial cell spheroids. Proteomics analysis of platelet-induced NETs in DR revealed dysregulation of proteins involved in platelet activation and NET formation, including CXCL7. Furthermore, increased CXCL-7 levels were observed in platelet pellet and plasma samples from the DR group. Additionally, CXCL7-treated neutrophils formed NETs via the CXCR2 receptor, and inhibition of NETosis was observed in neutrophils exposed to an anti-CXCL-7 antibody and a CXCR2 antagonist.

**Conclusion:** Our findings revealed that platelets released CXCL-7 induce NETs formation via the CXCL7/CXCR2 axis and blockade of CXCL7/CXCR2 axis inhibits the NETosis in DR, thereby inhibiting the pathogenesis of DR. Circulating CXCL7 serves as a potential prognostic marker, and the CXCL7/CXCR2 axis may be a therapeutic target for the treatment of DR.

**What Are the Clinical Implications?:** Platelets have emerged as immune cells, and platelet-neutrophil interactions are reported to play a significant role in the pathogenesis of various metabolic diseases. The role of platelet-neutrophil interactions and neutrophil extracellular traps (NETs) in the development of diabetic retinopathy (DR) remains poorly understood. Investigating the mechanistic regulators of platelet-mediated NETs in DR is crucial for identifying new therapeutic approaches. Our study observed increased platelet activation, platelet-neutrophil aggregates and NETs among DR subjects. Further, DR-platelet induces NET in healthy neutrophils, and NETs induce inflammation, angiogenesis, and disrupt endothelial barrier function in RF/6A cells in vitro. These findings strengthen the evidence that platelet-neutrophil interactions play a major role in DR pathogenesis. Proteomic analysis identified CXCL7 as a mechanistic regulator of platelet-induced NETs formation in DR. Inhibitors that target the platelet-derived CXCL7/CXCR2 axis for NETosis can be used for the prevention of retinal injury in DR. Overall, our work emphasises the mechanistic understanding of platelet-neutrophil interactions and CXCL7/CXCR2 axis as a therapeutic target for inhibiting the pathogenesis of DR.

**Graphical abstract:** Scheme for the platelet-derived CXCL7 regulation of NETs in DR. Activated platelets release CXCL7 and platelet aggregates with neutrophils to form NETs via the CXCL7:CXCR2 axis. Blockade of the CXCL7:CXCR2 axis by anti-CXCL7 antibody and CXCR2 inhibitor prevents NETs formation in DR.

## Introduction

Diabetic retinopathy (DR) is the most common microvascular complication of diabetes mellitus. It is characterised by chronic hyperglycaemia, systemic inflammation, endothelial dysfunction and neovascularisation [1]. DR is the leading cause of preventable blindness among working-age individuals. According to the 11th edition of the International Diabetes Federation (IDF) report, the prevalence of diabetic retinopathy among diabetic patients is 30%, with 10% of those experiencing vision-threatening diabetic retinopathy (VTDR) [2]. DR is associated with chronic systemic inflammation, which triggers immune responses through activation of immune cells [3]. In recent years, platelets have emerged as crucial immune cells that play a significant role in regulating inflammatory responses by interacting with various cells, including leukocytes and endothelial cells [4]. These interactions are mediated through the activation of platelet receptors (TLR-4, GPVI, Integrin αIIbβ3, P-selectin, and P2Y12) and the release of immune mediators, including PDGF, TGF-β, CXCL4 (PF4), CXCL8, RANTES (CCL5), and CXCL7 [5]. Among immune cells, neutrophils are the most abundant and serve as the primary responders to pathogens in innate immune system through various processes, including phagocytosis, degranulation, and the formation of neutrophil extracellular traps (NETs) [6]. NETs are web-like DNA structures formed by chromatin decondensation and extrusion from activated neutrophils. These NETs comprise cytotoxic granule proteins, including histones and serine proteases such as elastase, proteinase 3, and cathepsin G, which help eradicate pathogens, including viruses, bacteria, and fungi [7–9]. Deficient NET formation may facilitate pathogen immune evasion and exacerbate infections [10–11]. Additionally, NETs are formed in response to various pathological, physiological, and pharmacological stimuli, including cholesterol crystals, oxidised LDL, platelets, and platelet-released HMGB1 and cytokines [12]. Prolonged exposure to these stimuli can enhance NET formation and promote inflammation, resulting in tissue and organ damage [13–14]. NETs have a detrimental effect and are involved in the pathogenesis of various disease conditions, including autoimmune disorders, cardiometabolic diseases, vasculitis, atherosclerosis, thrombosis, stroke, and diabetic complications [15–16]. Wong et al. reported that prolonged diabetes primes neutrophils to form NETs, which interfere with wound healing in diabetic complications [17]. Additionally, increased levels of NETs in circulation, as well as NET deposits in the retina and vitreous fluid of patients with DR. This study also indicated that hyperglycemia induces the formation of NETs through reactive oxygen species (ROS) and NADPH oxidase [18]. Endothelial dysfunction is a characteristic feature of DR, and proteins associated with NETs, such as neutrophil elastase (NE), matrix metalloproteinase 9 (MMP9), and peptidylarginine deiminase 4 (PAD4), contribute to endothelial dysfunction and vascular damage [19]. However, the role of NETs in the pathogenesis of DR is less explored.

Besides the immunological factors, activated platelets aggregate with neutrophils to form platelet-neutrophil complexes (PNC), inducing NETs via TXA_2_, CXCL4 (PF4), vWF, GPIb and exacerbating the pathogenesis of various diseases, including transfusion-related acute lung injury (TRALI), ischemic stroke, and sepsis [20–22]. Inhibiting platelet activation and subsequent markers (CXCL4, TXA2, STING) prevented the NETs formation in disease conditions, highlighting the importance of platelet-mediated NETs as a potential therapeutic approach [23–25]. However, platelet-neutrophil interactions and their regulatory mechanisms, which contribute to the thromboinflamation in DR, are largely unexplored. Hence, studying the platelet-neutrophil interactions is crucial for identifying potential therapeutic targets and agents that aim to prevent or block increased platelet activation and the formation of NETs, which is vital for preventing or managing DR. Thus, in this study, we aimed to investigate the interaction between platelets and neutrophils, as well as the mechanistic regulators of platelet-mediated NETs in the development of DR.

## Materials and Methods

### Ethics statement, study participants and sample collection

The study was approved by the Institutional Ethics Committee of Sri Sankaradeva Nethralaya, Guwahati, with an Institutional Ethics Committee Approval No. SSDN/IEC/DEC/2021/06. Participants were recruited from the OPD of the vitreo-retina department at Sri Sankaradeva Nethralaya, Guwahati, after obtaining informed consent in accordance with the principles of the Declaration of Helsinki. Both male and female participants aged 30-70 years were included in the study. Participants with no prior history of T2DM, DR, or any other illness and who were not taking any medication were recruited as healthy controls (HC). Type 2 diabetes (T2DM) subjects were defined according to the ADA guidelines: prior history of T2DM, fasting blood glucose (FBG) ≥126 mg/dL, and glycosylated haemoglobin (HbA1c) ≥6.5%. DR subjects were enrolled according to the criteria for type 2 diabetes described above, and DR screening was performed by trained optometrists using a fundus photograph of both eyes (FF-450 fundus camera, ZEISS) after pupil dilation with tropicamide (1%). Based on the clinical features of the fundus images, two ophthalmologists confirmed the DR diagnosis and classified the NPDR and PDR cases according to the International Diabetic Retinopathy Disease Severity Scale proposed by Wilkinson et al. (2003) [26]. Subjects with eye conditions, including glaucoma, branch retinal vein occlusion (BRVO), central retinal vein occlusion (CRVO), macular hole, and age-related macular degeneration, as well as those with chronic conditions such as renal failure, liver failure, type 1 diabetes, thyroid disease, cancer or malignancy, from the study will be excluded. Clinical history and complete physical examination, including blood pressure, FBG (ACCU-CHECK Instant Glucose Meter and strips), and HbA1c (Accurex Xpress A1c Meter and strips), were obtained after informed consent from each participant. Peripheral blood samples were collected by venipuncture, using BD Vacutainer serum (SST) and plasma tubes (K2EDTA and ACD). Blood was allowed to clot by leaving it at room temperature for more than 15 minutes. After centrifugation at 1500g for 15 minutes at room temperature, the serum and plasma were aliquoted into 1.8 mL cryovials and immediately stored at - 80°C for further analysis. Blood biochemical profiles, including low-density lipoprotein (LDL), total cholesterol (TC), triglycerides (TG), high-density lipoproteins (HDL), AST, ALT, ALP, uric acid, urea, and creatinine levels, were measured using a biochemical analyser (Randox Dytona+, UK) according to standard protocols.

### Isolation of human platelets and neutrophils

Platelets were isolated from acid citrate dextrose (ACD)-anticogulated whole blood within 2 hrs of collection. Briefly, the whole blood was centrifuged at 200g for 20 min to separate platelet-rich plasma (PRP). Furthermore, PRP was mixed with HEPES buffer (1:1) and centrifuged at 100g for 10 minutes to separate the remaining red and white blood cells. The supernatant was centrifuged at 800g for 10 min to pellet the platelets. Platelet pellet washed three times with platelet wash buffer and resuspended in HEPES Tyrode’s buffer. Neutrophils were isolated from EDTA-anticogulated whole blood by BigSep Magnet (Stemcells, #18001) using EasySep Human Neutrophil Isolation Kit (Stemcells, #19666), with >95% viability observed in Thermo Countess 3.0 (Thermo Fisher Scientific).

### Flow cytometry

Platelet-rich plasma (PRP) was isolated from whole blood (BD Vacutainer® ACD tubes) by centrifugation at 500 rpm for 10 min at room temperature. Furthermore, 25 µL of PRP was incubated with an anti-human CD62P-APC conjugated antibody (BD, #550888) for 20 minutes at room temperature. Samples were washed and resuspended in 300 μL phosphate-buffered saline (PBS). Platelets were analysed using an Attune NxT flow cytometer (Thermo Fisher Scientific, Singapore). For NETs quantification, 100 μL of whole blood was lysed by a two-step procedure with RBC lysis buffer (GCC Biotech, #GRBC01), followed by washing with 400 μL of PBS and centrifugation at 400g for 10 min. Pellet was resuspended in 100 μL PBS and blocked with 10 μL anti-Hu Fc Receptor Binding Inhibitor Polyclonal Antibody (Thermo Fisher, #14-9161-73), followed by staining with anti-citruH3 (Citrulline R2+R8+R17) primary antibody (Abcam, #Ab5103) for 30 min. After washing with PBS, samples were further stained with anti-human CD66b-PE (BD Bioscience, #561650), anti-human CD41a-PE-Cy5 (BD Bioscience, #55968), anti-human MPO-FITC (BD Bioscience, #340580), and Goat anti-Rabbit, Alexa Fluor-700 Secondary Antibody (Thermo Fisher, #A21038), incubated for 30 mins at room temperature. Samples were washed and resuspended in 300 μL PBS and analysed by flow cytometry (Attune™ NxT Acoustic Focusing Cytometer). Granulocytes were gated on the FSC-SSC plot based on size and granularity. Further, CD66b-positive granulocytes were gated as neutrophils. Granulocytes positive for CD66b and CD41a were considered as platelet-neutrophil aggregate (PNA). Neutrophils (CD66b-positive granulocytes) positive for both CitruH3 and MPO were considered as NETotic neutrophils.

### Enzyme-linked immunosorbent assay (ELISA)

Human plasma CXCL4/PF4 (R&D systems, #DY-795), Neutrophil Elastase/ELA2 (R&D systems, #DY-9167-05), Proteinase 3/PRTN3 (R&D systems, #DY-6134), and Angiopoietin-2 (R&D systems, #DY-623) were measured by using ELISA kits as per the manufacturer’s protocol.

### Cell-free dsDNA quantification

Plasma Cell-free dsDNA levels were quantified using Quant-iT™ Pico Green® dsDNA assay Kit (Thermo, # P7589) as per the manufacturer’s protocol.

### Serum cytokine/chemokine, angiogenic markers and cell adhesion molecules measurement by multiplex analysis

Serum concentrations of cytokines/chemokines were measured using the Human ProcartaPlex Mix & Match 27-Plex (Invitrogen, #PPX-27-MX7DTFT) by the Bio-Plex 200 system (Bio-Rad Laboratories, Inc.). The customized panel includes Angiopoietin 1, CRP, Eotaxin (CCL11), E-selectin (CD62E), FGF-2, GM-CSF, ICAM-1, IFN gamma, IL-1 beta, IL-1RA, IL-2, IL-4, IL-8 (CXCL8), IL-10, IL-12p70, IL-13, IL-18, IP-10 (CXCL10), MCP-1 (CCL2), MIP-1 alpha (CCL3), MIP-1 beta (CCL4), PDGF-BB, PlGF-1, RANTES (CCL5), TNF alpha, VCAM-1, VEGF-A, VEGF-R2 (KDR). Serum samples stored at -80 °C were thawed on ice, vortexed and spun at 10000 x g for two minutes to remove debris. Each sample was analysed as per the manufacturer’s protocol. Data analysis was performed using Bio-Plex software (Version 7.1). Analyte concentrations were determined using 4 and 5 PL curve-fitting algorithms available in the software. Analytes (FGF-2, GM-CSF, IFN gamma, IL-1 beta, IL-1RA, IL-2, IL-4, IL-8 (CXCL8), IL-10, IL-12p70, IL-13, and PDGF-BB) falling out of the standard curve’s detection range were omitted for further analysis.

### Platelet neutrophil aggregation (PNA) and platelet-induced NETs formation in the co-culture assay

To determine the platelet-neutrophil aggregation (PNA) and platelet-induced NETs formation in DR, a platelet-neutrophil co-culture system was established. Briefly, neutrophils isolated from freshly collected whole blood of healthy participants at a density of 1×10^6^/ml were co-cultured with platelets (1×10^8^/ml) from HC, T2DM, and DR groups in poly-D-lysine (Thermo Fisher, # A38904-01) coated coverslips or cell culture plates (6 or 12-well plate) using RPMI1640 medium (10% FBS, 1% Anti-Anti, Gibco, #) at 37 °C and 5% CO2 for 6 hrs. To determine if CXCL7 induces NETs formation, neutrophils (1×10^6^/ml) from healthy participants were pre-treated with anti-CXCL7 antibody (R&D systems), CXCR2 inhibitor/SB225002 (TCI #182498-32-4) for 30 min followed by treatment with CXCL7 recombinant protein (200 ng/ml) in 6 or 12 well-plates using RPMI1640 medium (10% FBS) at 37 °C and 5% CO2 for 8 hrs.

### Isolation and quantification of NETs

Isolated neutrophils (1×10^6^cells /ml) were co-cultured with platelets (1×10^8^cells/ml) from HC, T2DM, and DR groups for 8 hrs, and with 400 nM of phorbol 12-myristate 13-acetate (PMA, Sigma, #P1585) for 3 hrs over poly-D lysine (Thermo Fisher, # A38904-01) coated coverslips and culture plates. After removing the supernatant, the deposited layer of NETs in the culture dish was washed with PBS and treated with PBS containing 10U/mL of DNase I (Sigma, #10104159001) in the presence of CaCl_2_ (2mM) at 37°C for 20 minutes, followed by the addition of EDTA to a final concentration of 10 mM. NETs were collected by pipetting vigorously. To remove cell debris, samples were centrifuged at 3000 x g for 10 minutes, and supernatants containing NETs were used for further experiments. The dsDNA content was measured using the PicoGreen dsDNA kit (Invitrogen, #P7589), according to the manufacturer’s protocol.

### Immunofluorescence assay

To visualise the PNA, platelets were stained for F-actin with AF-488 Phalloidin, and neutrophils were counterstained with DAPI nuclear dye. Platelet-neutrophil interactions were visualised using LEICA SP8 laser confocal microscopy (63x oil-immersed lens). For platelet-induced NETs, slides were stained with anti-human MPO (Thermo Fisher, #A11OO1) and anti-CD61 (Thermo Fisher, #MA5-32077) antibodies, followed by secondary antibodies anti-mouse AF-488 and anti-rabbit AF-700. The slides were then counterstained and mounted using ProLong Gold antifade reagent with DAPI (Invitrogen, #P36941). Slides were visualised using LEICA STELARIS WLL confocal microscopy (63x oil-immersed lens).

### Cell culture and in vitro assays using RF/6A cell line

The RF/6A cell line is a monkey choroidal–retinal vascular endothelial cell line, and it was generously gifted by Dr Prajakta Dandekar Jain’s lab (Institute of Chemical Technology, Mumbai). Cells were cultured in Minimum Essential Medium (MEM), supplemented with 10% fetal bovine serum (FBS) and 1% antibiotic-antimycotic solution (Anti-Anti, Gibco). The cells were cultured in a humidified incubator at 37 °C with 5% CO2. To determine the effect of NETs on RF/6 cells, cells were seeded in either a six or 12-well plate in MEM (10% FBS) media. After achieving 90-100% confluency, cells were treated with 600 ng/ml and 900 ng/ml DNA conc. of NETs for 24 hrs.

### Sprouting assay

RF/6A cells were trypsinised at 80% to 90% confluency, and spheroids were formed using the hanging-drop method as previously reported, with some modifications [27]. Spheroids were embedded in the collagen I matrix in a 96-well plate ultra-low attachment surface (Corning, #3474) at 2-3 spheroids/well. After polymerisation (30 min at 37 °C), spheroids were treated with basal media, NETs (600, 900 ng/ml) and VEGF-A (50 nM, Abclonal, #RP01162) as a positive control for 24 hrs at 37 °C and 5% CO2. After 24 hrs, the spheroids were fixed using 10% PFA for 2 hrs. Spheroids were stained for F-actin using AF-488 Phalloidin dye, and nuclei were counter-stained using DAPI (2 μg/ml). Spheroids were imaged using a Nikon Eclipse Ti2 fluorescence microscope using a 10x objective lens. Images were further analysed using the FIJI ImageJ software (JAVA 21.0.7).

### Trans-endothelial electrical resistance (TEER) assay

To evaluate the integrity of the RF/6A endothelial cell monolayer, we conducted TEER assay using epithelial voltmeter (EVOM2, World Precision Instruments). The RF/6A cells were cultured in a 12-well plate cell culture insert (pore size 0.4 μm, surface area 1.13 cm^²^, Thermo Fisher Scientific, #140652) for 14 to 21 days, until a monolayer formed. The monolayers were treated with NETs (600 ng/ml and 900 ng/ml) for 24 hrs. The TEER (Ω/cm²) for each cell culture insert was calculated by subtracting the resistance of a blank cell culture insert (Ω, media without cells) from the measured resistance of each cell culture insert (Ω) and normalising to the membrane surface area (cm^²^) of each cell culture insert.

### Immunoblotting

To determine the levels of CitruH3, CXCL7, and ZO-1 in NETs, platelets, and NETs-treated RF/6A cells, respectively, total protein was extracted using T-PER reagent (Thermo Scientific, #78510) containing 1x EDTA solution (Thermo Scientific, #1861283) and 1x Halt Protease & Phosphatase Inhibitor Cocktail (Thermo Scientific, #1861284). Protein concentrations were quantified using the micro-BCA kit (Thermo Scientific, #23235) following the manufacturer’s protocol. 40 μg of total protein was loaded in 8-15% SDS-PAGE and transferred to the PVDF membrane (0.45 μm pore size, Millipore, #IPVH00010). Blots were blocked with 5% BSA for 1 hour at room temperature, followed by three washes with TBST. Further blots were incubated overnight in citruH3, CXCL7 and ZO-1 primary antibody solution (1:1000) in 2-3% BSA solution at 4 °C. After washing three times with TBST solution, blots were incubated with secondary antibody solution for 1 hr, followed by rinsing with TBST solution three times. The blots were developed using West Pico/Femto substrate solution (Thermo Scientific, #34577 & #34095), and images were captured using Fusion S (Vilber Lourmat, Collégien, France). Images were quantified using ImageJ (version 21.0.7).

### RNA isolation and quantitative real-time PCR analysis

Total RNA was isolated from the RF/6A cells using Trizol reagent (Thermo Fisher Scientific, #15596026). The concentration of the isolated RNA was determined using a microplate reader (Epoch BioTek, USA). Complementary DNA (cDNA) was synthesised by reverse transcription with the ABscript II cDNA First Strand Synthesis Kit (ABclonal, #RK20400), following the manufacturer’s instructions. Gene-specific primers for the target genes were mixed with the synthesised cDNA and subjected to quantitative real-time PCR using the QuantStudio 5 Real-Time PCR System (Applied Biosystems # A34322). The reaction mixture included cDNA, forward and reverse primers, TB Green Premix Ex Taq, and nuclease-free water. The thermal cycling conditions for each gene were as follows: an initial denaturation at 95°C for 5 minutes; 40 cycles of 95°C for 15 seconds, annealing temp. of 52-58°C for 30 seconds (Table S1), and 72°C for 30 seconds; followed by a final extension at 72°C for 5 minutes and a hold at 4°C for 2 minutes. Relative gene expression was calculated using the 2^-ΔΔCt method, with GAPDH serving as the internal control for normalisation (StepOne Plus Software V.). The primer sequences and their respective annealing temperatures are provided in **Table S1** in the **Supplementary file.**

### Proteomics analysis of platelet-induced NETs

#### In vitro NETs induction using platelets and sample preparation for proteomic analysis

To identify the mechanistic regulators of platelet-mediated NETs in the context of DR pathogenesis, we conducted a comprehensive analysis of proteomic profiles of NETs formed by platelets from HC, T2DM, and DR (n=4 from each group) using quantitative (relative) Swath-MS analysis. Briefly, Isolated neutrophils (1×10^6^/ml) from the healthy group were incubated with unstimulated platelets (1×10^8^/ml) from each study group in RPMI-1640 medium (10% FBS) at 37°C and 5% CO_2_ for 8 hrs. After incubation, the supernatant was carefully removed without disturbing the deposited NETs. The NETs were then washed in PBS and incubated with PBS containing 10 U/ml DNase I (Sigma, #10104159001) and CaCl2 (2 mM) for 20 min at 37°C. DNase activity was halted by adding EDTA to a final concentration of 10 mM. The samples were collected using vigorous pipetting and then centrifuged at 3000 × g for 10 min to remove cell debris. The supernatants containing the NETs were used for proteomics sample preparation, and a previously published method was used for proteomic analysis with some modifications [28]. Detailed trypsin digestion and SWATH-MS analysis were mentioned in the **Supplementary file.**

### Statistical analysis

Normalised data are presented as mean ± standard deviation, while non-normalised data are shown with median and interquartile range (Q1-Q3). The difference between the study groups was tested using One-way ANOVA, followed by Tukey’s multiple comparison tests, and Kruskal-Wallis, followed by Dunn’s test, to assess the statistical significance between different groups. P<0.05 is considered statistically significant. Statistical analysis was performed using PRISM, version 7.0 (GraphPad) and R software (version 25.05.01+513). The heatmap was visualised using the pheatmap package. Spearman correlation analysis and visualisation were performed using the ggcorrplot and ggplot packages. Differential protein expression analysis was carried out to identify proteins with significant changes using log_10_FC <-0.18 and log_10_FC >0.18 and p value <0.05 as cutoff for significantly downregulated and upregulated proteins, respectively. The DEPs were plotted in a volcano plot to visually represent the distribution of these proteins. Gene Ontology (GO) enrichment analysis was performed to identify functional biological terms associated with upregulated and downregulated proteins. The analysis was conducted using the Enrichr web tool (https://maayanlab.cloud/Enrichr/), for upregulated and downregulated proteins separately across three GO categories: biological process, cellular component, and molecular function. Statistical significance was determined based on Benjamini-Hochberg corrected p-values (adjusted p-value) ≤ 0.05. Each GO category with an adjusted p-value ≤ 0.05 was included. The plot was created using the ggplot2 and ggalluvial packages to visualize the gene ontologies.

## Results

### Subject characteristics

The study included participants from four groups: healthy controls (HC), individuals with type 2 diabetes mellitus(T2DM), those with non-proliferative DR (NPDR), and individuals with proliferative DR (PDR). The detailed clinical and biochemical characteristics of the study groups are presented in Table 1.

**Table 1:**
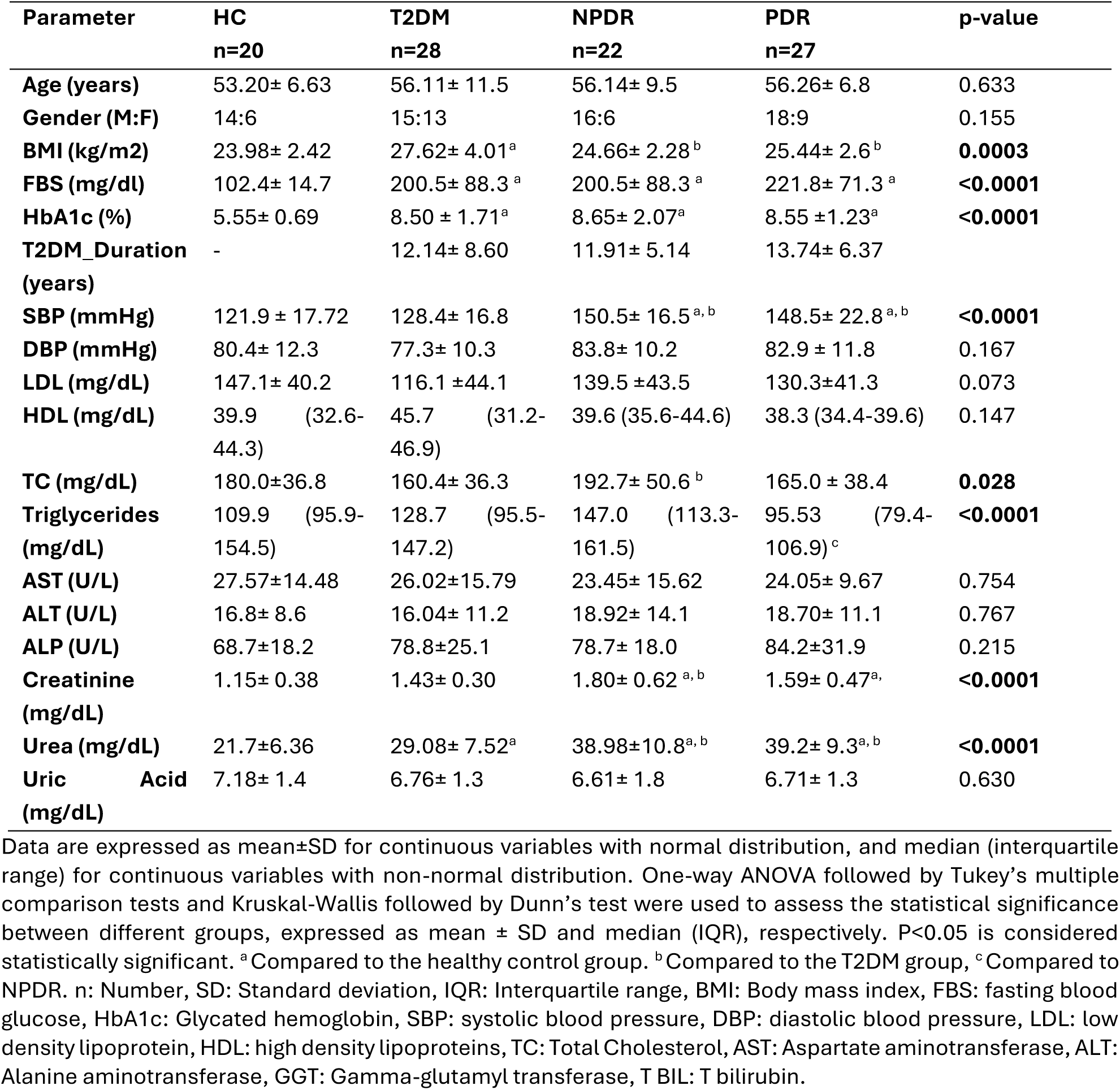
Clinical and biochemical characteristics of subjects.

### Increased platelet activation is associated with NETs formation in diabetic retinopathy

To evaluate the role of platelet activation and platelet-neutrophil interaction in the progression of DR, we quantified platelet activation by measuring p-selectin expression of the platelets, platelet-neutrophil aggregates (PNA), and % of NETotic neutrophils and % of NETs among platelet-neutrophil aggregates and non-aggregating platelets using flow cytometry, in clinical samples of study groups. Our results showed a significant and progressive increase in platelet activation p-selectin (%) expression on platelets and plasma levels of platelet factor 4 (PF4/CXCL4) in patients with T2DM and DR compared to HC participants (Fig. 1C-1D). Moreover, we observed a significant and progressive increase in % of platelet-neutrophil aggregates (PNA) among T2DM and DR participants (Fig. 1E), and these findings suggest that activated platelets form aggregates with neutrophils in DR. Additionally, % of NETotic neutrophils was significantly elevated in DR patients compared to HC and T2DM patients (Fig. 1F). The study also revealed that NETotic neutrophils were significantly higher among the platelet neutrophil aggregates (PNA) than non-aggregating neutrophils (Fig. 1G). Furthermore, the circulatory NETs marker, i.e., plasma dsDNA, was significantly higher in DR (both NPDR and PDR) patients (Fig. 1J). We found that plasma neutrophil elastase (NE) levels were significantly higher only in the advanced stage of DR (PDR) compared to other study groups (Fig. 1K), and a gradual significant increase in plasma proteinase 3 (PR3) levels was observed in subjects with DR subjects compared to HC and T2DM subjects (Fig. 1L). Additionally, we found a significant positive correlation between platelet activation markers (p-selectin and PF4 levels) and the NETs markers (%NETs, dsDNA, NE, Fig. 1M). These findings suggest that activated platelets form aggregates with neutrophils, which leads to the formation of NETs in patients with DR.

**Figure 1:**
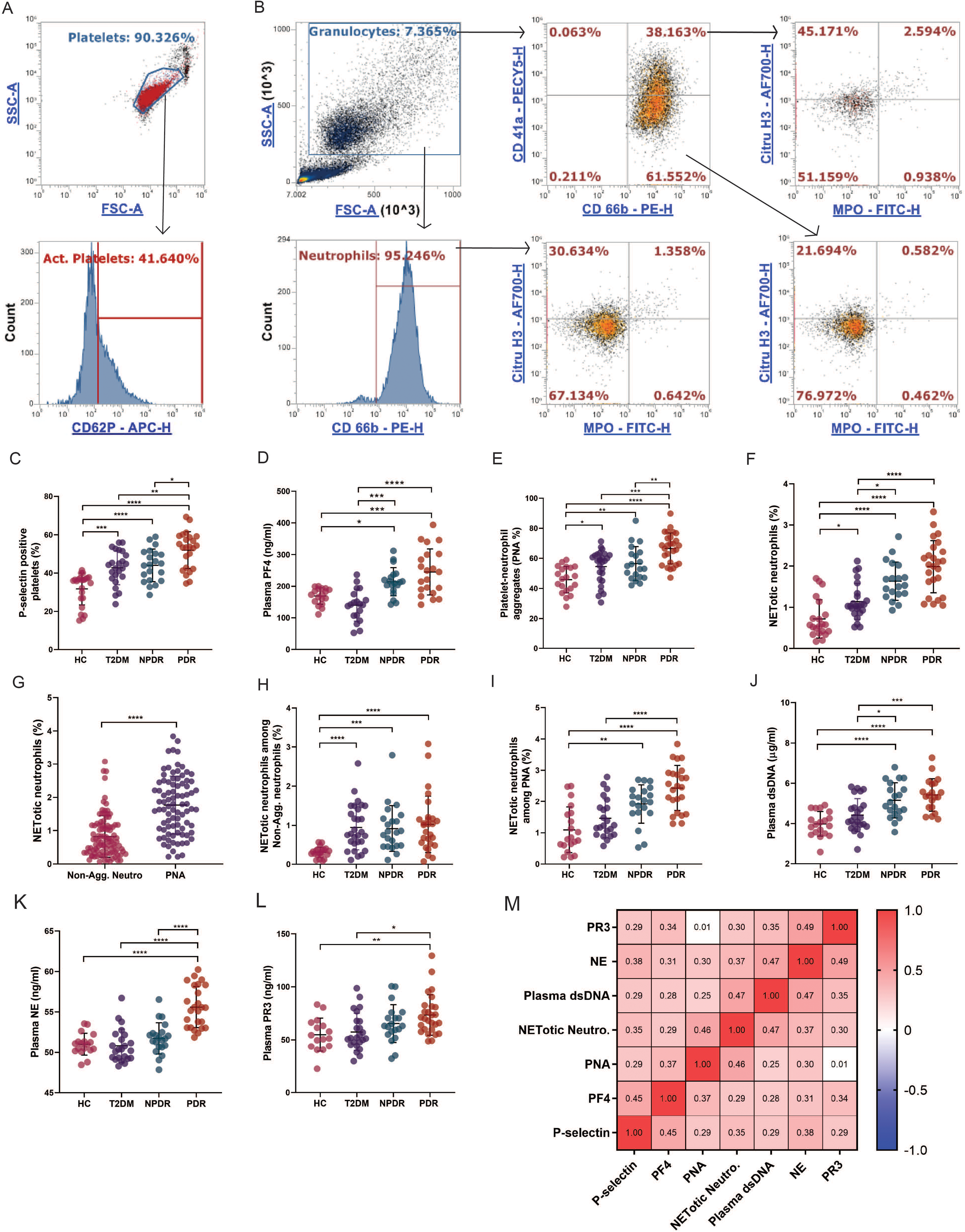
Increased platelet activation, PNA and NETs markers in DR. **A)** Gating strategy for platelet activation; platelets were gated based on forward and side scatter characteristics from platelet-rich plasma. Among platelets, events positive for CD62p (P-selectin) were gated as activated platelets. **B)** Gating strategy for platelet-neutrophil aggregates (PNA) and NETs; Granulocytes were gated using forward and side scatter characteristics from RBC-lysed whole blood. Further, CD66b-positive granulocytes were gated as neutrophils. Among neutrophils, events positive for CD41a were considered as platelet-neutrophil aggregates (PNA). Events positive for both MPO and citrullinated histone-3 among neutrophils, PNA and non-aggregating neutrophils were considered as NETotic neutrophils. Quantification of **C)** P-selectin. **D)** plasma PF4. **E)** % of platelet-neutrophil aggregates (PNA). **F)** % of NETotic neutrophils among study groups (HC, T2DM, NPDR, and PDR). **G)** % of NETotic neutrophils among non-aggregating neutrophils and platelet-neutrophil aggregates. **H)** % of NETotic neutrophils among non-aggregating neutrophils in study groups. **I)** NETotic neutrophils among platelet-neutrophil aggregates in study groups. **J)** plasma cell-free dsDNA. **K)** plasma neutrophil elastase (NE). **L)** plasma proteinase 3 (PR3), among study groups. **M)** Spearman correlation matrix of platelet activation, PNA and NETosis markers. p< 0.05 is considered significant. *p < 0.05, **p < 0.01, ***p < 0.001 and ****p<0.0001. (HC: n=20, T2DM: n=28, NPDR: n=22, PDR: n=28), Abbreviations: HC= healthy Control, T2DM= type 2 diabetes mellitus, NPDR= non-proliferative diabetic retinopathy, PDR= proliferative diabetic retinopathy, PNA= platelet-neutrophil aggregates, NETs= neutrophil extracellular traps, PF4= platelet factor 4, NE= neutrophil elastase, PR3= proteinase 3, MPO=myeloperoxidase, citruH3= citrullinated histone 3.

### DR platelets aggregate with neutrophils and form NETs in vitro

To further confirm the role of platelet-neutrophil interactions in DR, we co-incubated platelets from HC, T2DM and DR with healthy neutrophils (1:200 ratio) for 4 hrs. Our observations revealed that DR platelets aggregated with healthy neutrophils compared to the platelets from T2DM and HC subjects **(Fig. 2A).** Additionally, we examined the role of DR platelets in the formation of NETs. In this study, platelets from the HC, T2DM, and DR groups were co-incubated with healthy neutrophils (at a 1:200 ratio) for 8 hrs and observed DR platelets induced the formation of NETs in healthy neutrophils, while no NETs were observed in neutrophils co-cultured with the T2DM and HC platelets **(Fig. 2B).** To further validate these findings, we visualized samples using scanning electron microscopy (SEM), which revealed the presence of PNA and NET-like structures in neutrophils co-cultured with DR platelets. Although T2DM platelets did form aggregates with healthy neutrophils, NET-like structures were not detected **(Fig. 2C).** Moreover, we conducted western blot analysis for citrullinated H3 protein (NETs protein) using lysates from the platelet-neutrophil co-cultures. The analysis demonstrated increased citrullination of H3 in samples from DR platelets co-cultured with neutrophils **(Fig. 2D).** These results confirm our hypothesis that platelet-neutrophil interactions drive NET formation and are involved in the pathogenesis of DR.

**Figure 2:**
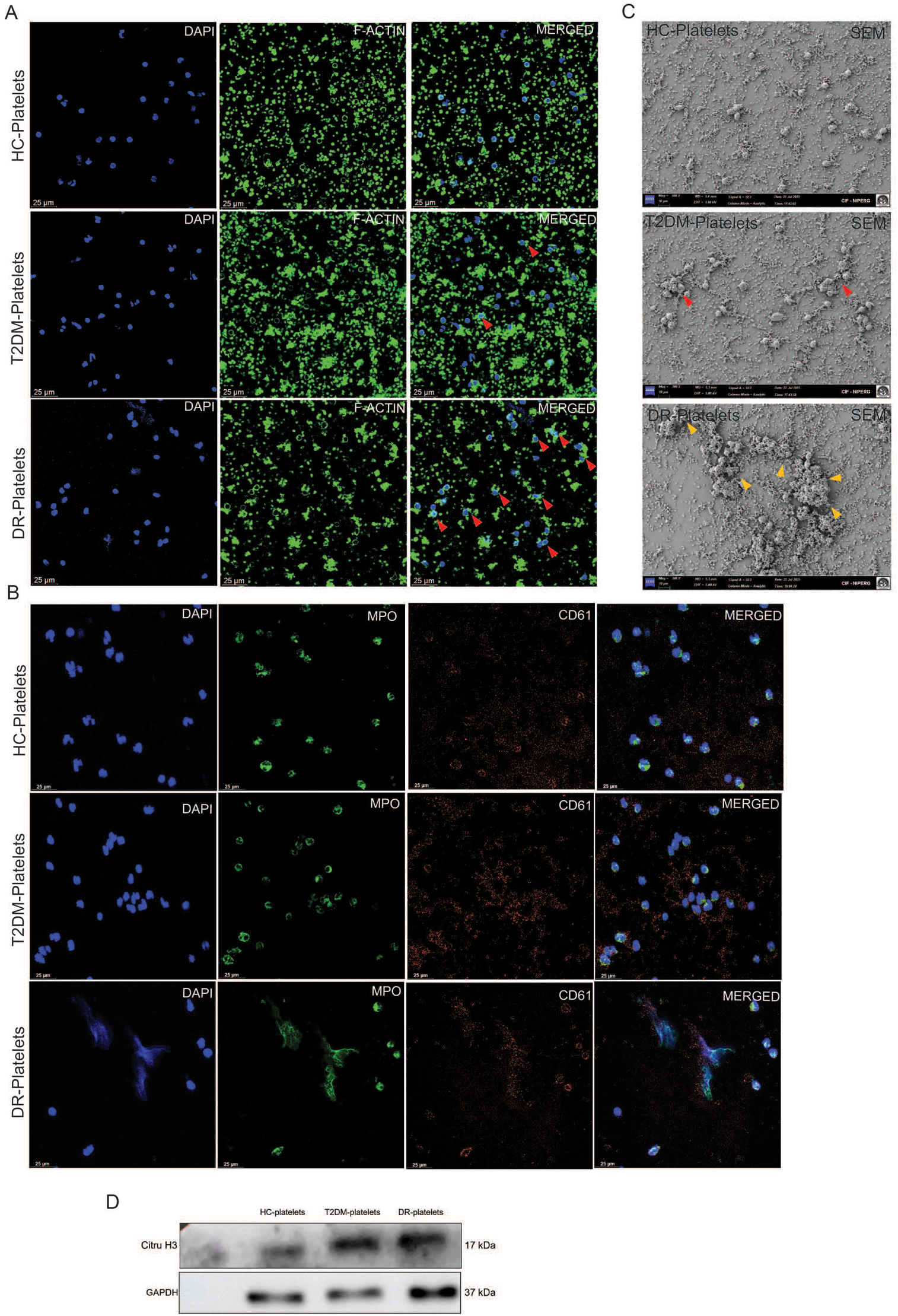
DR platelets aggregate with neutrophils and form NETs in vitro. **A)** Neutrophils (1×10^6^/ml) were incubated with platelets (1×10^8^/ml) from HC, T2DM and DR patients on Poly-D lysine coated coverslips for 4 hours. The cells were fixed and stained with phalloidin-AF488 (F-actin, green) and DAPI (nucleus, blue). The slides were imaged using a Leica SP8 LASER confocal microscope (63x magnification, size bar 25 μm). DR platelets formed aggregates with neutrophils (red arrows). **B-C)** Neutrophils (1×106/ml) were incubated with platelets (1×108/ml) from HC, T2DM and DR patients on Poly-D lysine coated coverslips for 8 hours. **B)** The cells were fixed and immunostained for MPO (NETs, green) and CD61 (platelets, red), followed by DAPI ( DNA staining, blue). The slides were visualised using Leica SP8 LASER confocal microscope (63x magnification, size bar 25 μm). DR platelets induced NETs formation in neutrophils. **C)** The cells were fixed and dried by ethanol fractional dehydration for scanning electron microscopy (SEM). SEM images also show DR platelets aggregated with neutrophils and formed NETs structures (yellow arrows). **D)** Protein levels of Citru H3 were assessed by western blotting.

### Platelet activation and NETs markers association with systemic inflammation, angiogenesis and endothelial dysfunction in diabetic retinopathy

To investigate the relationship between activated platelets and NETs and systemic inflammation, endothelial dysfunction, and angiogenesis, we analysed circulating levels of inflammatory cytokines, chemokines, endothelial dysfunction markers, and angiogenic markers. The heatmap illustrates relative changes **(Fig. 3A)** and median log2 fold changes (with respect to HC; **Fig. 3B) in** circulatory inflammatory cytokines/chemokines and angiogenic markers across the study groups. Among the inflammatory cytokines/chemokines, IL-18 and RANTES were significantly increased in DR subjects compared to healthy subjects and IP-10 levels were significantly reduced in both T2DM and DR subjects. The study found no significant differences in CRP, MCP1, PlGF1, and eotaxin levels among the study groups. Regarding endothelial dysfunction and angiogenic markers, the circulatory levels of pro-angiogenic markers such as VEGF-A and ANGPT2 were significantly elevated in DR patients. Additionally, the circulatory level of endothelial dysfunction marker ICAM-1 was elevated in DR (NPDR and PDR) patients. Increased levels of E-selectin were observed only in patients with NPDR, indicating its involvement in the early stage of DR. The anti-angiogenic marker ANGPT1 was significantly decreased in both T2DM and DR patients compared to healthy controls. These findings underscore the pro-inflammatory state and endothelial dysfunction observed in patients with DR. A Spearman correlation analysis was performed to explore potential relationships among platelet activation and NET markers, circulating cytokines/chemokines, and angiogenic markers. In this study, IL-18 exhibited a significant positive correlation with platelet activation and NETotic neutrophils. Similarly, VEGFA and ANGPT2 showed positive correlations with platelet activation and markers of PNA and NETs (NETotic neutrophils, dsDNA, NE, and PR3). Conversely, ANGPT1 and IP-10 showed a significant inverse correlation with platelet activation, PNA, and NETs markers (NETotic neutrophils, dsDNA, and NE) (Fig. 3C). These data indicate a significant relationship between circulatory cytokines/chemokines and angiogenic markers, on the one hand, and platelet activation, PNA, and NETs markers, on the other.

**Figure 3:**
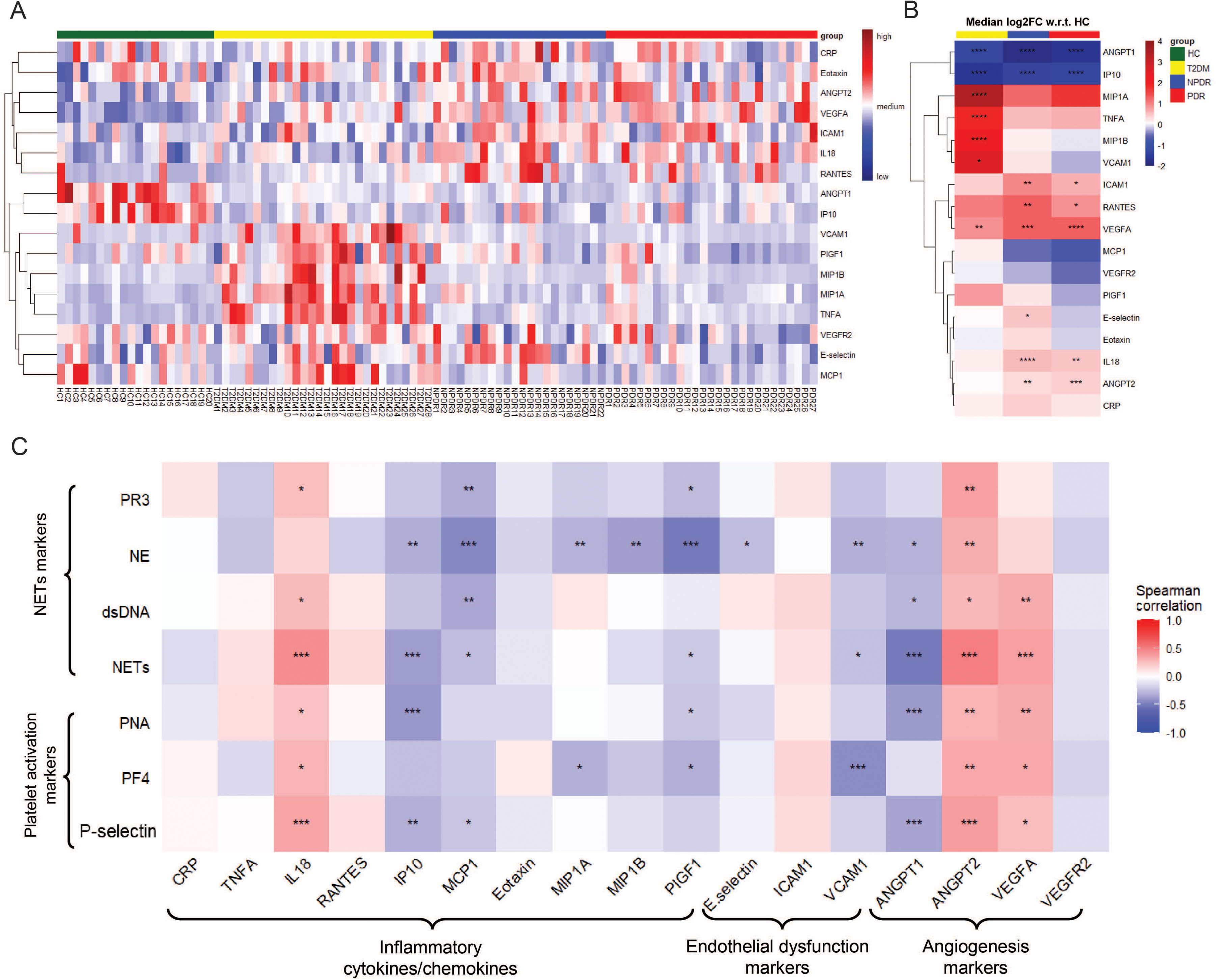
Platelet activation markers, platelet-neutrophil aggregates (PNA) and NETs were associated with inflammation, endothelial dysfunction and angiogenesis in DR. **A)** Heat map represents relative quantification of the inflammatory cytokines/chemokines, endothelial dysfunction and angiogenesis markers among HC, T2DM, NPDR and PDR groups. **B)** Heatmap represents median log2 fold change of inflammatory cytokines/chemokines and angiogenic markers among T2DM, NPDR and PDR groups with respect to the HC. **C)** Spearman correlational matrix representing correlation between platelet activation, PNA, NETs markers and inflammatory cytokines/chemokines, endothelial dysfunction and angiogenic markers. p< 0.05 is considered significant. *p < 0.05, **p < 0.01, ***p < 0.001 and ****p<0.0001.

### NETs exacerbate endothelial dysfunction and inflammation in choroid-retinal endothelial cells in vitro

Endothelial dysfunction and inflammation are characteristic features of the DR. This study investigated the effects of NETs on inflammation and endothelial dysfunction in retinal endothelial cells (RF/6A). The mRNA expression of inflammatory markers, i.e., TNF-alpha, IL-1 beta, IL-6, and IL-8, and endothelial dysfunction markers, i.e., ICAM1 and VCAM1, was quantified in RF/6A cells treated with NETs. **Fig. 4A and 4B** illustrate the relative mRNA expression of these markers across the study groups. The results showed elevated mRNA expression of inflammatory cytokines, including IL-1β, IL-6, and IL-8, upon NETs treatment. Additionally, markers of endothelial dysfunction, including ICAM1, VCAM1 and E-selectin, were also found to be elevated after NETs exposure. Our results indicate that NETs exposure exacerbates the endothelial inflammation and endothelial injury in the retina.

**Figure 4:**
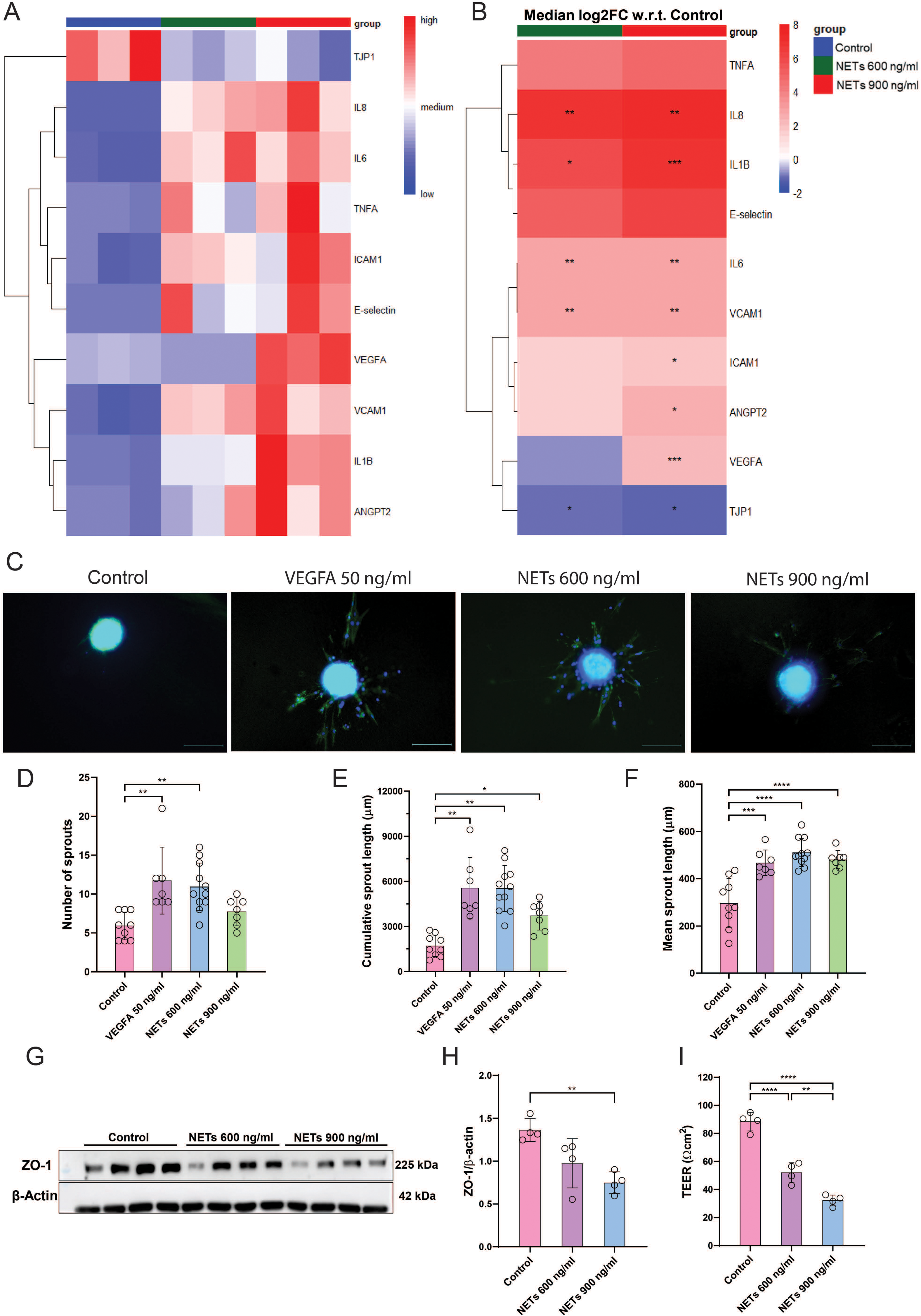
NETs induced inflammation, endothelial dysfunction, angiogenesis and endothelial barrier disruption in RF/6A cells in vitro. RF/6A cells were exposed to 600 ng/ml and 900 ng/ml concentrations of NETs for 24 hours. **A)** Heat map represents relative quantification of the mRNA expression of inflammatory markers, endothelial dysfunction, and angiogenic markers (n=3 for each group). **B)** Heatmap represents median log2 fold change of serum inflammatory cytokines/chemokines and angiogenic markers with respect to the HC (n=3 for each group). **C)** RF/6A spheroids embedded in collagen matrices in 96-well ULA plates were exposed to NETs at 600 ng/ml and 900 ng/ml concentrations for 24 hours. Spheroids were fixed and stained with phalloidin-AF488 (F-actin, green) and DAPI (nucleus, blue). The spheroids were imaged using Nikon ECLIPSE Ti2 fluorescent microscope (10x magnification, size bar 500 μm). **D-E)** Quantification of protein levels of ZO-1 in RF/6A cells on NETs exposure by western blotting (n=4 for each group). **F.** Effect of NETs on RF/6A cell monolayers was assessed by TEER assay. RF/6A cell monolayer cultured in 12-well plate cell culture inserts (0.4 μm pore size, 1.33 cm^2^ surface area) exposed to NETs at 600 ng/ml and 900 ng/ml concentrations for 24 hours, and TEER was measured by EVOM^2^ (World Precision Instruments) (n=4 for each group). p< 0.05 was considered significant. *p < 0.05, **p < 0.01, ***p < 0.001 and ****p<0.0001.

### NETs promote angiogenesis in choroid-retinal endothelial cells in vitro

To investigate the effect of NETs on the angiogenesis of retinal RF/6A cells, we treated collagen matrix-embedded spheroids of RF/6A cells with different concentrations of NETs (600 ng/ml and 900ng/ml). The number of sprouts, average sprout length, cumulative sprout length, and number of branches per sprout were quantified for the treatment groups. The data indicate that spheroids treated with NETs had significantly higher numbers of sprouts, greater cumulative sprout length, and longer mean sprout length compared to untreated spheroids **(Fig. 4C-F)**. Consistently, we observed mRNA overexpression of proangiogenic genes, including VEGFA and ANGPT2, in RF/6A cells after NETs exposure (Fig. 4A). This indicates that NETs exacerbate angiogenesis in retinal endothelial cells and contribute to neovascularisation in DR.

### NETs impair the integrity of the retinal endothelial cell barrier in vitro

We further investigated the impact of NETs on the endothelial monolayer barrier, as the integrity of the endothelial barrier relies on the junctional proteins in the monolayer. To evaluate the effect of NETs on ZO-1 (TJP-1) in the RF/6A monolayer using western blotting techniques. This study observed a significant reduction in ZO-1 expression at both the protein and mRNA levels following NETs treatment compared to the control group **(Fig. 4A-B and 4G-H)**. Additionally, we performed a TEER assay to assess the integrity of the endothelial cell monolayer (Transwells Thermo Fisher Scientific, #140652). Results revealed a significant decrease in electrical resistance in the monolayer after 24 hrs of NETs treatment compared to the control group **(Fig. 4I).** These findings demonstrate the detrimental effect of NETs on the endothelial barrier. Therefore, NETs break the retinal endothelial barrier, facilitating the leakage of blood components in the vitreous compartment in DR.

### Proteomic analysis reveals the potential mediators of platelet-induced NETs in diabetic retinopathy

The current findings of the study highlight that activated platelets drive the formation of NETs in patients with DR. To investigate the molecular mechanism of platelet-induced NETs in DR patients, we performed proteomics analysis of platelet-induced NET samples **(Fig. 5A)**. The analysis identified 339 proteins. Among them, 193 proteins were common across all three groups (HC, T2DM, and DR), while 20 proteins were common in HC-platelets and T2DM-platelets induced NETs. Additionally, 78 proteins were common to HC-platelets and DR-platelets that induced NETs, and only three were common to T2DM-platelets and DR-platelets that induced NETs. Furthermore, 31 proteins were unique to T2DM-platelets, and 13 were unique to DR-platelets that induced NETs **(Fig. S1)**. The differently expressed proteins (DEPs) were plotted in a volcano plot to visually represent their distribution. Various platelet-related proteins (WDR-1, vWF-1, CXCL7) were differentially expressed in DR-platelet groups as compared to the T2DM-platelet and HC-platelet groups. NETs-related proteins, i.e., NE (ElANE), PERM, NAGL, Histone H3 & H4, DEF3 were identified in all study groups. Differential expression analysis was carried out to identify proteins with significant changes in their expression levels using cutoffs of log_10_ fold change <-0.18 (for downregulated proteins) and >0.18 (for upregulated proteins), along with p < 0.05 **(Fig. 5B)**.

**Figure 5:**
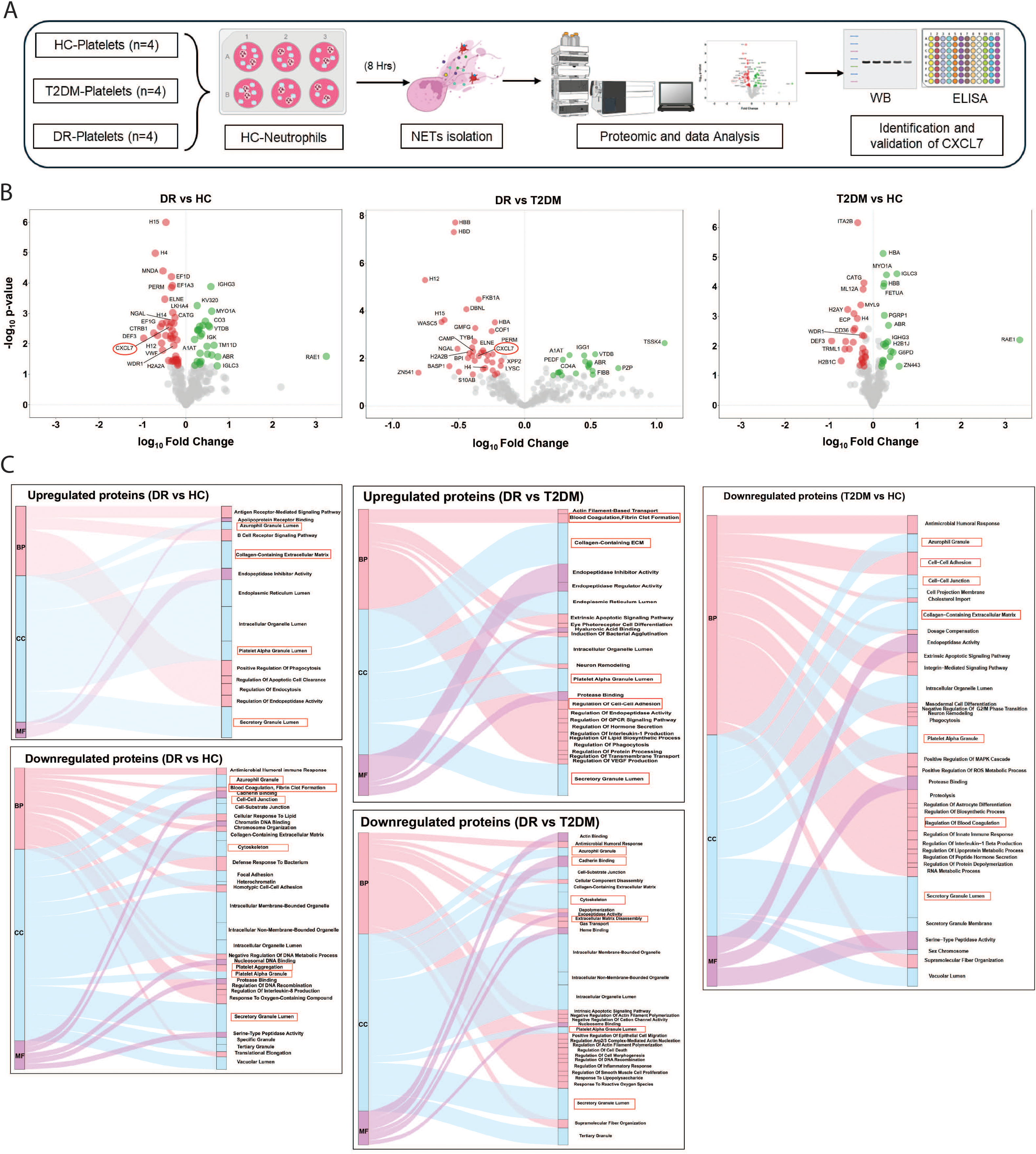
Platelet-derived CXCL7 is a mediator for NETs formation. **A)** In vitro study design for proteomics analysis of Platelet-induced NETs. **B)** Volcano plot represents the differentially expressed proteins among platelet-induced NETs for DR vs HC, T2DM vs HC, and DR vs T2DM. **C)** Gene ontology terms among upregulated and downregulated proteins for different comparison groups (n=4).

Furthermore, Gene Ontology (GO) enrichment analysis identified functional biological terms associated with the differentially expressed proteins, for upregulated and downregulated proteins. Regarding cellular components (CC), the differentially expressed proteins in DR-platelets were associated with the cytoskeleton, the azurophil granule lumen, and the platelet alpha granule lumen. The enriched biological processes (BPs) for differentially expressed proteins included antimicrobial humoral immune response, antigen-receptor-mediated signalling pathways, blood coagulation, regulation of IL-8 production, and regulation of cell-cell adhesion **(Fig. 5C)**. Furthermore, we explored CXCL7 as a potential mediator of platelet-induced NETs in DR.

### Platelet-derived CXCL7 as a key mediator for inducing NETosis and a predictive marker for diabetic retinopathy

CXCL7 is a protein found in platelets and is known to be a potent chemoattractant for neutrophils at the sites of injury [29]. Previous studies reported that CXCL7 is involved in the pathophysiology of various diseases; its role in inducing NETosis in diabetic retinopathy has not yet been explored. In our study, we quantified the levels of CXCL7 in both platelets and circulation among study groups (HC, T2DM, & DR). Western blot analysis of CXCL7 in platelet lysate from different study groups revealed that platelets from the DR group exhibited significantly elevated CXCL7 expression (normalised to Beta-actin) compared to both HC and patients with T2DM (Fig. 6A-B). Similarly, plasma CXCL7 levels were also significantly elevated in the DR group compared to T2DM and HC groups **(Fig. 6C)**. Further, we explored the potential relationship between plasma CXCL7 and platelet activation markers (P-selectin & PF4), platelet-neutrophil aggregates (PNA), and NETs markers (% of Netotic neutrophils, dsDNA, NE, and PR3). Spearman correlation analysis revealed a significant positive correlation between plasma CXCL7 and platelet activation markers, specifically p-selectin and PF4; furthermore, it was positively correlated with PNA, % of NETs, and plasma dsDNA levels (**Fig. 6D-J)**. Furthermore, CXCL7 demonstrated potential as a diagnostic marker for differentiating between DR vs HC, DR vs T2DM, and DR vs non-DR (combined HC & T2DM) with an area under the curve (AUC) of 0.80, 0.87, and 0.84, respectively (**Fig. 6K-M)**. These results confirm that CXCL7 is the key mediator for NETs formation involved in the pathophysiology of diabetic retinopathy, and CXCL7 is a predictive marker for diabetic retinopathy.

**Figure 6:**
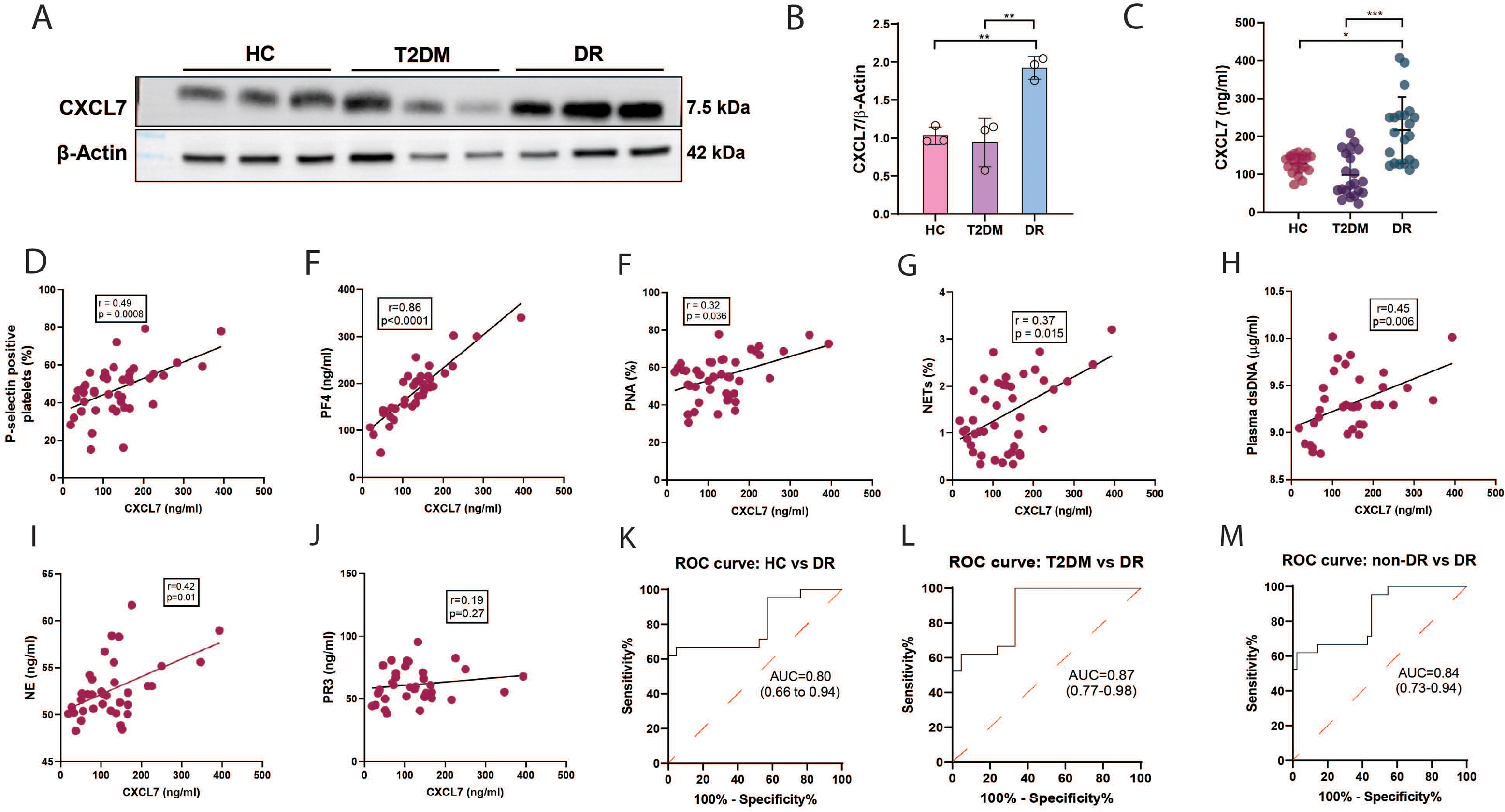
Quantification and validation of CXCL7. **A-B)** Protein levels of CXCL7 in platelet lysate were quantified by western blotting (n=4). **C)** Quantification of plasma levels of CXCL7 among HC, T2DM, and DR groups using ELISA assay (n=21 each group). Spearman correlation of plasma CXCL7 with **D)** % of p-selectin positive platelets**. E)** plasma levels of PF4**. F)** percentage of platelet-neutrophil aggregates**, G)** percentage of NETotic neutrophils. **H)** plasma levels of dsDNA. **I)** plasma levels of NE. **J)** plasma levels of PR3. ROC curve for CXCL7 for discriminating between **K)** HC and DR**. L)** T2DM and DR, and **M)** non-DR and DR. Control and T2DM subjects will be included as non-DR. p< 0.05 was considered significant. *p < 0.05, **p < 0.01, and ***p < 0.001.

### Blockade of the CXCL7:CXCR2 axis inhibits the NETs formation in vitro

Our data suggest a positive relationship between circulatory CXCL7 levels and platelet activation, as well as NET markers in clinical samples. We further investigated the role of CXCL7 in the formation of NETs and explored the potential of blocking CXCL7 as a therapeutic target for inhibiting NETs formation. Our flow cytometry data revealed that neutrophils treated with CXCL7 (200 ng/mL) showed a significantly higher percentage of NETotic neutrophils than untreated healthy neutrophils. CXCL7 is a potent ligand for the CXCR2 receptor, and blockade of CXCR2 with a CXCR2 antagonist (SB255002) significantly reduced the NETotic neutrophils (**Fig. 7A-C).** Similarly, western blot data show the significant overexpression of Citru H3 protein in CXCL7-exposed neutrophils as compared to healthy neutrophils. The CXCR2 antagonist significantly reduced the Citru H3 expression **(Fig. 7D-E).** These results were corroborated by immunofluorescence analysis, which revealed the presence of NETs among neutrophils treated with CXCL7, in contrast to those in the control group. Additionally, a significant reduction in NETs formation was observed when CXCL7 and CXCR2 were blocked by an anti-CXCL7 antibody (2ug/ml) and CXCR2 antagonist (SB255002), respectively **(Fig. 7F).** These findings suggest that CXCL7 induces NETs formation via the CXCR2 receptor and blocking the CXCL7:CXCR2 axis can prevent NETs formation, emphasising the therapeutic potential of targeting the CXCL7:CXCR2 axis to prevent NETs formation and the subsequent pathogenesis of DR.

**Figure 7:**
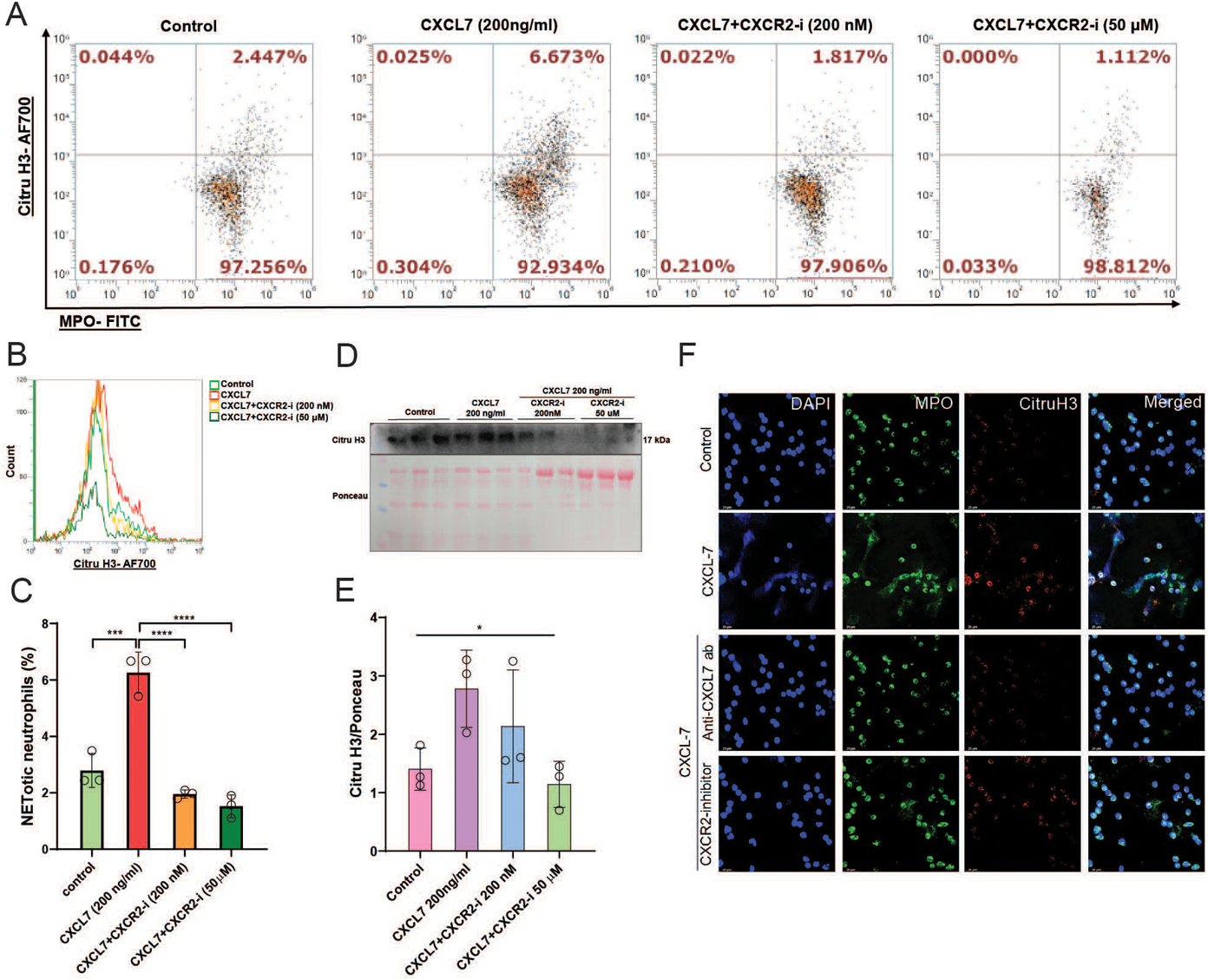
CXCL7 induces NETs formation, and blockade of the CXCL7: CXCR2 axis inhibits the NETs formation in vitro. **A-C)** Flow cytometry data showing the NETs formation among control, CXCL7 (200 ng/ml), and CXCL7 (200 ng/ml) with CXCR2 inhibitor (200nM & 50 μM) treated neutrophils **C)** CXCL7 (200 ng/ml) significantly induced NETs among neutrophils compared to control. CXCL7 (200 ng/ml) did not induce NETs in neutrophils pretreated with CXCR2 inhibitor (200nM & 50 Μm) (n=3). **D-E)** Protein levels of Citru H3 were assessed by western blotting. Similarly, CXCL7 (200 ng/ml) induced overexpression of citru H3 among neutrophils compared to control. Expression of citru H3 was reduced in CXCL7 (200 ng/ml) treated neutrophils pretreated with CXCR2 inhibitor (200nM & 50 μM) (n=3). **F)** Immunofluorescence imaging also shows that CXCL7 (200 ng/ml) induces NETs among neutrophils compared to control. CXCL7 (200 ng/ml) did not induce NETs when neutralised with anti-CXCL7 antibody (2 μg/ml) and in neutrophils pretreated with CXCR2 inhibitor (50 μM) (Leica SP8 LASER confocal microscope, 63x magnification, size bar 25 μm). p< 0.05 was considered significant. *p < 0.05, **p < 0.01, ***p < 0.001 and ****p<0.0001.

## Discussion

Despite the established role of NETosis in DR, the interaction between platelets and neutrophils, as well as the mechanisms regulating platelet-induced NETs in DR, remains largely unexplored. In this study, first, we observed a significant increase in platelet activation, PNA, and NETs in DR, and platelet-mediated NETs were confirmed *in vitro*. Second, platelet activation and NETs markers were associated with circulating inflammatory cytokines/chemokines, as well as angiogenesis markers in DR. Third, upon exposure to NETs, retinal endothelial cells (RF/6A) exhibited pro-inflammatory and pro-angiogenic phenotypes, along with disruption of the endothelial barrier. Fourth, proteomic analysis of platelet-induced NETs, along with further validation, identified platelet-derived CXCL7 as a potential marker for NET formation in DR. CXCL7 alone induced NETs *in vitro*, and inhibition of either CXCL7 or its receptor, CXCR2, prevented NET formation. Recognising the critical role of platelet-induced NET formation and its mechanistic regulators in diabetic retinopathy marks a significant shift that could lead to novel therapeutic strategies.

Recently, platelets have been identified as immune cells that interact with other immune cells, playing a crucial role in the inflammatory response. Upon activation, platelets release proteins from dense alpha granules, including cytokines/chemokines, growth factors, and metabolites [4–5]. Platelet activation is associated with chronic inflammation of diabetes and its complications, including coronary artery diseases (CAD), diabetic nephropathy (DN), and DR [30–31]. In the present study, we observed increased levels of the platelet activation markers p-selectin and PF4 in DR patients. Similarly, we also observed elevated PNA among DR patients. Activated platelets aggregate with neutrophils through p-selectin/PSGL-1 and αMβ2 integrin/glycoprotein Ibα (GPIbα) interactions, leading to the formation of platelet-neutrophil complexes and facilitating the adhesion and transmigration of neutrophils through the endothelial cells [32]. Furthermore, we observed elevated levels of NETs and NET markers, such as dsDNA and NE, in DR, suggesting NET involvement in the pathogenesis of DR. This finding is consistent with previous studies [18, 33].

Additionally, the extent of NETs was more pronounced among PNA compared to non-aggregating neutrophils. In line with these findings, an in vitro investigation confirmed that DR platelets aggregated with healthy neutrophils and induced NET formation. Furthermore, platelet activation markers are associated with the percentage of NETs and NETosis markers. These findings indicate the critical role of platelet-neutrophil interactions in NETs formation in DR. It is reported that platelets activated by various agonists, including TRAP, LPS, ADP, and thrombin, were able to aggregate with neutrophils and induce NETs formation through distinct mechanisms, including the interaction of p-selectin with PSGL-1, activation of HMGB-1 by TLR4 and lipid peroxidation [34–36]

In chronic hyperglycaemia, platelet-neutrophil interactions drive inflammation by inducing the production of cytokines and chemokines [32]. In our study, we observed increased levels of IL-18 and CCL5 in the DR group. IL-18, a marker of NLRP3 inflammasome activation, was elevated in DR and showed a positive correlation with platelet activation and NETs markers. NLRP3 inflammasome activation plays a critical role in chronic inflammation, promotes platelet pyroptosis, induces NET formation, and retinal neovascularisation [37–39]. Consistently, we observed overexpression of IL-1β, an inflammasome activation-dependent marker, in RF/6A cells after NET exposure. This highlights the role of inflammasome activation in platelet-neutrophil interactions and endothelial inflammation in DR. CCL-5 is a potential systemic platelet-released marker, and increased serum CCL-5 enhances retinal vascular permeability, monocyte/macrophage trafficking and inflammatory response in retinal tissues in DR [40–41]. Furthermore, endothelial dysfunction is a characteristic feature of DR, characterised by overexpression of adhesion molecules, including E-selectin, ICAM1, and VCAM1, which facilitate the attachment and transmigration of leukocytes through the endothelium [42].

Similarly, a significant increase in serum ICAM-1 and E-selectin levels was observed in the DR group. Additionally, RF/6A cells showed mRNA overexpression of ICAM-1 and VCAM-1 upon exposure to NETs, consistent with previous literature [43–45]. These findings highlight the positive feedback between NET formation and endothelial dysfunction. Similarly, VEGFA, ANGPT1, and ANGPT2 are growth factors that regulate endothelial barrier integrity in DR [46–47]. As in previous findings, VEGFA and ANGPT2 were elevated in DR, whereas ANGPT1 was significantly decreased [46–47]. We observed a positive correlation between platelet activation and NETs with pro-angiogenic markers, including VEGFA and ANGPT2, highlighting their involvement in platelet activation and NETs in angiogenesis. Upon further investigation, an in vitro study revealed that NETs induced the mRNA overexpression of pro-angiogenic markers (ANGPT-2, VEGFA) and promoted angiogenesis, as evidenced by the sprouting assay in endothelial cells. Similarly, NETs have been reported to induce proangiogenic effects on endothelial cells previously [48].

The involvement of platelets in inducing NETs formation was first reported in severe sepsis, later in transfusion-related acute lung injury (TRALI), SLE, myocardial infarction (MI), deep vein thrombosis, pulmonary thrombosis and stroke [20, 49–51]. However, platelet-neutrophil interactions are not well explored in DR. In the current study, a proteomics investigation of platelet-induced NETs identified proteins involved in platelet activation (CXCL7, vWF, WDR1), platelet-neutrophil interactions, and NET formation. Activated platelets release various chemokines and cytokines from their granules, which serve as powerful mediators of immune defence and inflammatory processes. Some of these platelet-derived agents include the C-X-C motif chemokines CXCL4 (PF4), CXCL7, CXCL11, CXCL12, and CXCL14 [52]. CXCL7/NAP-2 is predominantly a platelet protein and is reported to be a potent chemoattractant for neutrophils and other immune cells, exhibiting pro-inflammatory and pro-angiogenic effects [29]. CXCL7 plays a significant role in the pathogenesis of various inflammatory diseases, including its importance in NETosis under different conditions, such as cancer and atrial fibrillation [53–54]. However, the role of CXCL-7 in the pathogenesis of DR remains unexplored. Therefore, among the differentially expressed proteins, we selected CXCL7 to evaluate its role in regulating NETs. To identify the role of CXCL-7 in DR pathogenesis in DR patients, increased CXCL7/NAP2 expression was observed in platelet lysate and plasma in DR. Plasma CXCL7 concentrations showed an association with platelet activation markers such as p-selectin and PF4, as well as with PNAs and NETs markers, including NETs, neutrophil elastase (NE), and proteinase 3 (PR3). These findings suggest a strong link between CXCL7 and both platelet activation and NETs formation in DR. Supporting evidence from animal studies has demonstrated that elevated CXCL7 levels in the retinas of DR rat models reinforce its potential role in disease pathology [55]. Similarly, earlier clinical work by F.E. Preston and colleagues reported increased β-thromboglobulin, a precursor of CXCL7, in diabetic patients with microangiopathy, which correlated positively with platelet aggregation [56]. Together, these observations highlight CXCL7 as a key mediator linking platelet activation, NETs, and microvascular complications in DR. CXCL7 is a ligand for both CXCR1 and CXCR2 receptors; however, it is more specific for CXCR2 than for CXCR1 [53]. Various studies have reported the involvement of the CXCL7:CXCR1/2 axis in thrombo-inflammation and endothelial dysfunction in various diseases, including cerebral aneurysm and renal carcinoma [57–58]. Blockade of CXCR1/2 has been shown to significantly inhibit thrombosis and lung injury in sepsis by inhibiting NET formation [59].

Inhibiting or blocking the platelet-neutrophil interaction and the formation of NETs, as well as modulating NETs-related markers, reduces the pathogenesis of various diseases, including atherosclerosis, inflammatory bowel disease (IBD), and sepsis [60–62]. Targeting platelet-neutrophil interactions by blocking PF4 prevented thrombosis in deep vein thrombosis (DVT) [63]. Similarly, blockade of HMGB1 reduced infarct volume and improved neurological deficits in photothrombotic stroke (PTS) by inhibiting platelet-induced NET formation [60]. Furthermore, PAD4 inhibitors reduced NET formation and, subsequently, endothelial integrity during atherosclerotic plaque formation [61]. Targeting NET-related proteins, such as NE, MPO, MMP9, citrullinated histone 3, and dsDNA, alleviated NET formation. Targeting dsDNA using DNase I neutralised the NETs in IBD [62]. Excessive NET formation is associated with tissue injuries in sepsis, and NE inhibitor Sivelestat alleviates the tissue injury during septic shock [64]. Similarly, in our study, neutrophils pretreated with anti-CXCL7 antibody and CXCR2 inhibitor did not form NETs in the presence of CXCL7. These results highlight the role of the CXCL7:CXCR2 axis in NETs formation in DR. Similarly, Mohmad Alsabani et al. 2021 highlighted that blockade of CXCR1/2 receptors inhibits NETs formation in sepsis patients [59]. Blockade of the NET formation by targeting the CXCL7/CXCR2 axis can inhibit the detrimental effects of NETs, including endothelial dysfunction, angiogenesis and disruption of the endothelial barrier in diabetic retinopathy.

## Conclusion

In summary, the prolonged diabetic conditions activate platelets. Activated platelets release CXCL7, which attracts the neutrophils toward the platelets, forming the platelet-neutrophil complexes, which induce CXCL7 release and promote the formation of NETs by binding to the CXCR2 receptor. NETs further induce endothelial dysfunction and disrupt the retinal endothelial cell barrier. Blockade of the CXCL7:CXCR2 axis using an anti-CXCL7 antibody and CXCR2 inhibitor (SB225002) significantly inhibited NET formation. Therefore, the CXCL7:CXCR2 axis could be exploited to inhibit NET formation in DR. Inhibitors that target the platelet-derived CXCL7-CXCR2 NETosis axis can be used for the prevention of retinal injury in DR.

## Declarations

### Ethics approval and consent to participate

The study was approved by the Institutional Ethics Committee of Sri Sankaradeva Nethralaya, Guwahati, with an Institutional Ethics Committee Approval No. SSDN/IEC/DEC/2021/06. Participants were recruited from the OPD of the vitreo-retina department of Sri Sankaradeva Nethralaya, Guwahati, after taking informed consent per the principles of the Declaration of Helsinki.

### Consent for publication

Not applicable

### Availability of data and materials

All data will be shared by the corresponding author upon reasonable request.

### Authors’ contributions

BN and RA have designed the whole study. BN has performed most of the experiments and SM was involved in other in vitro experiments. BN and MJB have recruited study patients, collected clinical data and performed related assays. MJK have performed the SWATH-based proteomic analysis. RS and AKY performed proteomics data analysis. AG and SS helped with Western blotting. AA performed ELISA quantification. BN and RA have performed the final analysis and interpretation of the data. RA was responsible for overseeing the project and preparing the final manuscript. All authors approved the final version of the manuscript. RA is responsible for the integrity of the work as a whole.

## Acknowledgements

We would like to thank all study participants for their generosity and commitment, without whom this work would not have been possible. We thank Director NIPER Guwahati and Founder Director and President Dr. Harsha Bhattacharjee for their support. Dr. Prasenjit Guchait for his valuable suggestions during the study. Dr. Prajakta Dandekar Jain for her generosity in providing RF/6A cell line.

## Supplementary Information

The online version of this article contains peer-reviewed but unedited supplementary file.

## References

[1] Wong TY, Cheung CMG, Larsen M, et al. Diabetic retinopathy. Nat Rev Dis Primers 2016;2:16012. doi: 10.1038/nrdp.2016.12.

[2] International Diabetes Federation. IDF Diabetes Atlas, 11th edn. Brussels: International Diabetes Federation, 2025. (available from https://diabetesatlas.org).

[3] Zhao B, Zhao Y, Sun X. Mechanism and therapeutic targets of circulating immune cells in diabetic retinopathy. Pharmacol Res 2024;210:107505. doi: 10.1016/j.phrs.2024.107505.

[4] Morrell CN, Aggrey AA, Chapman LM et al. Emerging roles for platelets as immune and inflammatory cells. Blood 2014;123:2759–2767. doi: 10.1182/blood-2013-11-462432.

[5] Tokarz-Deptuła B, Baraniecki Ł, Palma J et al. Characterization of platelet receptors and their involvement in immune activation of these cells. Int J Mol Sci 2024;25:12611. doi: 10.3390/ijms252312611

[6] Gierlikowska B, Stachura A, Gierlikowski W et al. Phagocytosis, degranulation and extracellular trap release by neutrophils: the current knowledge, pharmacological modulation and future prospects. Front Pharmacol 2021;12:666732. doi: 10.3389/fphar.2021.666732.

[7] Brinkmann V, Reichard U, Goosmann C et al. Neutrophil extracellular traps kill bacteria. Science 2004;303:1532–1535. doi: 10.1126/science.1092385.

[8] Fuchs TA, Abed U, Goosmann C et al. Novel cell death program leads to neutrophil extracellular traps. J Cell Biol 2007;176:231–241. doi: 10.1083/jcb.200606027.

[9] Korba-Mikołajczyk A, Służalska KD, Kasperkiewicz P. Exploring the involvement of serine proteases in neutrophil extracellular traps: a review of mechanisms and implications. Cell Death Dis 2025;16:535. doi: 10.1038/s41419-025-07857-w.

[10] Baz AA, Hao H, Lan S et al. Neutrophil extracellular traps in bacterial infections and evasion strategies. Front Immunol 2024;15:1357967. doi: 10.3389/fimmu.2024.1357967.

[11] Meier A, Sakoulas G, Nizet V et al. Neutrophil extracellular traps: an emerging therapeutic target to improve infectious disease outcomes. J Infect Dis 2024;230:514–521. doi: 10.1093/infdis/jiae252.

[12] Gao F, Peng H, Gou R et al. Exploring neutrophil extracellular traps: mechanisms of immune regulation and future therapeutic potential. Exp Hematol Oncol 2025;14:80. doi: 10.1186/s40164-025-00670-3.

[13] Zuo Y, Yalavarthi S, Shi H et al. Neutrophil extracellular traps in COVID-19. JCI Insight 2020;5:e138999. doi: 10.1172/jci.insight.138999.

[14] Retter A, Singer M, Annane D. ‘The NET effect’: neutrophil extracellular traps: a potential key component of the dysregulated host immune response in sepsis. Crit Care 2025;29:59. doi: 10.1186/s13054-025-05283-0.

[15] Papayannopoulos V Neutrophil extracellular traps in immunity and disease. Nat Rev Immunol 2018;18:134–147. 10.1038/nri.2017.105.

[16] Njeim R, Azar WS, Fares AH et al. NETosis contributes to the pathogenesis of diabetes and its complications. J Mol Endocrinol 2020;65:R65–R76. doi: 10.1530/jme-20-0128.

[17] Wong SL, Demers M, Martinod K et al. Diabetes primes neutrophils to undergo NETosis, which impairs wound healing. Nat Med 2015;21:815–819. doi: 10.1038/nm.3887.

[18] Wang L, Zhou X, Yin Y et al. Hyperglycaemia induces neutrophil extracellular traps formation through an NADPH oxidase-dependent pathway in diabetic retinopathy. Front Immunol 2019;9:3076. doi: 10.3389/fimmu.2018.03076.

[19] Yu S, Liu J, Yan N. Endothelial dysfunction induced by extracellular neutrophil traps plays important role in the occurrence and treatment of extracellular neutrophil traps-related disease. Int J Mol Sci 2022;23:5626. doi: 10.3390/ijms23105626.

[20] Caudrillier A, Kessenbrock K, Gilliss BM et al. Platelets induce neutrophil extracellular traps in transfusion-related acute lung injury. J Clin Invest 2012;122:2661–2671. doi: 10.1172/JCI61303.

[21] Gao X, Zhao X, Li J et al. Neutrophil extracellular traps mediated by platelet microvesicles promote thrombosis and brain injury in acute ischemic stroke. Cell Commun Signal 2024;22:50. doi: 10.1186/s12964-023-01379-8

[22] Yang M, Jiang H, Ding C et al. STING activation in platelets aggravates septic thrombosis by enhancing platelet activation and granule secretion. Immunity 2023;56:1013–1026. doi: 10.1016/j.immuni.2023.02.015.

[23] Matsumoto K, Yasuoka H, Yoshimoto K et al. Platelet CXCL4 mediates neutrophil extracellular traps formation in ANCA-associated vasculitis. Sci Rep 2021;11:222. doi: 10.1038/s41598-020-80685-4.

[24] Carestia A, Kaufman T, Rivadeneyra L et al. Mediators and molecular pathways involved in the regulation of neutrophil extracellular trap formation mediated by activated platelets. J Leukoc Biol 2016;99:153–162. doi: 10.1189/jlb.3a0415-161r.

[25] Mizurini DM, Aslan JS, Gomes T et al. Salivary thromboxane A2-binding proteins from triatomine vectors of Chagas disease inhibit platelet-mediated neutrophil extracellular traps (NETs) formation and arterial thrombosis. PLoS Negl Trop Dis 2015;9:e0003869. doi: 10.1371/journal.pntd.0003869.

[26] Wilkinson CP, Ferris FL III, Klein RE et al. Global Diabetic Retinopathy Project Group. Proposed international clinical diabetic retinopathy and diabetic macular edema disease severity scales. Ophthalmology 2003;110:1677–1682. doi: 10.1016/S0161-6420(03)00475-5.

[27] Tetzlaff F, Fischer A. Human endothelial cell spheroid-based sprouting angiogenesis assay in collagen. Bio Protoc 2018;8:e2995. doi: 10.21769/BioProtoc.2995.

[28] Bai S, Chaurasiya AH, Banarjee R, et al. MJ. CD44, a predominant protein in methylglyoxal-induced secretome of muscle cells, is elevated in diabetic plasma. ACS Omega 2020;5:25016–25028. doi: 10.1021/acsomega.0c01318.

[29] Baggiolini M, Walz A, Kunkel SL. Neutrophil-activating peptide-1/interleukin 8, a novel cytokine that activates neutrophils. J Clin Invest 1989;84:1045–1049. doi: 10.1172/JCI114265.

[30] Kong YX, Rehan R, Moreno CL et al. SEC61B regulates calcium flux and platelet hyperreactivity in diabetes. J Clin Invest 2025;135:e184597. doi: 10.1172/JCI184597.

[31] Ferroni P, Basili S, Falco A et al. Platelet activation in type 2 diabetes mellitus. J Thromb Haemost 2004;2:1282–1291. doi: 10.1111/j.1538-7836.2004.00836.x.

[32] Shrimali NM, Agarwal S, Tiwari A et al. Platelet–neutrophil interactions and thrombo-inflammatory complications in type 2 diabetes mellitus. Curr Pathobiol Rep 2022;10:1–10. doi: 10.1007/s40139-022-00229-5.

[33] Binet F, Cagnone G, Crespo-Garcia S et al. Neutrophil extracellular traps target senescent vasculature for tissue remodeling in retinopathy. Science 2020;369:eaay5356. doi: 10.1126/science.aay5356.

[34] Zucoloto AZ, Jenne CN. Platelet–neutrophil interplay: insights into neutrophil extracellular trap (NET)-driven coagulation in infection. Front Cardiovasc Med 2019;6:85. doi: 10.3389/fcvm.2019.00085.

[35] Hoppenbrouwers T, Autar AS, Sultan AR et al. In vitro induction of NETosis: comprehensive live imaging comparison and systematic review. PLoS One 2017;12:e0176472. doi: 10.1371/journal.pone.0176472.

[36] Ono M, Toyomoto M, Yamauchi M et al. Platelets accelerate lipid peroxidation and induce pathogenic neutrophil extracellular trap release. Cell Chem Biol 2024;31:2085–2095. doi: 10.1016/j.chembiol.2024.11.003.

[37] Paik S, Kim JK, Shin HJ et al. Updated insights into the molecular networks for NLRP3 inflammasome activation. Cell Mol Immunol 2025;22:1–34. doi: 10.1038/s41423-025-01284-9.

[38] Su M, Chen C, Li S et al. Gasdermin D-dependent platelet pyroptosis exacerbates NET formation and inflammation in severe sepsis. Nat Cardiovasc Res 2022;1:732–747. doi: 10.1038/s44161-022-00108-7.

[39] Zhao P, Zhu J, Bai L et al. Neutrophil extracellular traps induce pyroptosis of pulmonary microvascular endothelial cells by activating the NLRP3 inflammasome. Clin Exp Immunol 2024;217:89–98. doi: 10.1093/cei/uxae028.

[40] Machlus KR, Johnson KE, Kulenthirarajan R et al. CCL5 derived from platelets increases megakaryocyte proplatelet formation. Blood 2016;127:921–926. doi: 10.1182/blood-2015-05-644583.

[41] Meleth AD, Agrón E, Chan CC et al. Serum inflammatory markers in diabetic retinopathy. Invest Ophthalmol Vis Sci 2005;46:4295–4301. doi: 10.1167/iovs.04-1057.

[42] Simó-Servat O, Simó R, Hernández C. Circulating biomarkers of diabetic retinopathy: an overview based on physiopathology. J Diabetes Res 2016;2016:5263798. doi: 10.1155/2016/5263798.

[43] Folco EJ, Mawson TL, Vromman A et al. Neutrophil extracellular traps induce endothelial cell activation and tissue factor production through interleukin-1α and cathepsin G. Arterioscler Thromb Vasc Biol 2018;38:1901–1912. doi: 10.1161/ATVBAHA.118.311150.

[44] Lin S, Zhu P, Jiang L et al. Neutrophil extracellular traps induced by IL-1β promote endothelial dysfunction and aggravate limb ischemia. Hypertens Res 2024;47:1654–1667. doi: 10.1038/s41440-024-01661-3.

[45] Wu S, Chen Y, Jin X et al. Toll-like receptors promote high glucose-induced vascular endothelial cell dysfunction by regulating neutrophil extracellular traps formation. Inflammation 2025;48:1–8. doi: 10.1007/s10753-025-02283-8.

[46] Shakthiya T, Chand L, Annamalai R et. Exploring the association of VEGF-A and ANGPTL2 with the prognosis of non-proliferative and proliferative diabetic retinopathy. Cureus 2024;16:e68273. doi: 10.7759/cureus.68273.

[47] Wang Y, Fang J, Niu T et al. Serum Ang-1/Ang-2 ratio may be a promising biomarker for evaluating severity of diabetic retinopathy. Graefes Arch Clin Exp Ophthalmol 2023;261:49–55. doi: 10.1007/s00417-022-05745-z.

[48] Aldabbous L, Abdul-Salam V, McKinnon T, Duluc L, Pepke-Zaba J, Southwood M, Ainscough AJ, Hadinnapola C, Wilkins MR, Toshner M, Wojciak-Stothard B. Neutrophil extracellular traps promote angiogenesis: evidence from vascular pathology in pulmonary hypertension. Arterioscler Thromb Vasc Biol 2016;36:2078–2087. doi: 10.1161/atvbaha.116.307634.

[49] Clark SR, Ma AC, Tavener SA et al. Platelet TLR4 activates neutrophil extracellular traps to ensnare bacteria in septic blood. Nat Med 2007;13:463–469. doi: 10.1038/nm1565.

[50] Maugeri N, Campana L, Gavina M et al. Activated platelets present high mobility group box 1 to neutrophils, inducing autophagy and promoting the extrusion of neutrophil extracellular traps. J Thromb Haemost 2014;12:2074–2088. doi: 10.1111/jth.12710.

[51] Stakos DA, Kambas K et al. Expression of functional tissue factor by neutrophil extracellular traps in culprit artery of acute myocardial infarction. Eur Heart J 2015;36:1405–1414. doi: 10.1093/eurheartj/ehv007.

[52] Flad HD, Brandt E. Platelet-derived chemokines: pathophysiology and therapeutic aspects. Cell Mol Life Sci 2010;67:2363–2386. doi: 10.1007/s00018-010-0306-x.

[53] Wu Q, Tu H, Li J. Multifaceted roles of chemokine CXC motif ligand 7 in inflammatory diseases and cancer. Front Pharmacol 2022;13:914730. doi: 10.3389/fphar.2022.914730.

[54] Ząbczyk M, Natorska J, Matusik PT et al. Neutrophil-activating peptide 2 as a novel modulator of fibrin clot properties in patients with atrial fibrillation. Transl Stroke Res 2024;15:773–783. doi: 10.1007/s12975-023-01169-6.

[55] Augustine J, Troendle EP, Friedel T et al. Scavenging acrolein with 2-HDP preserves neurovascular integrity in a rat model of diabetic retinal disease. Diabetologia 2025;68:2609–2629. doi: 10.1007/s00125-025-06515-2.

[56] Preston FA, Ward JD, Marcola B et al. Elevated β-thromboglobulin levels and circulating platelet aggregates in diabetic microangiopathy. Lancet 1978;311:238–240. doi: 10.1016/S0140-6736(78)90557-7.

[57] Nowicki KW, Mittal A, Hudson JS et al. Blockade of the platelet-driven CXCL7–CXCR1/2 inflammatory axis prevents murine cerebral aneurysm formation and rupture. Transl Stroke Res 2025;16:1251–1261. doi: 10.1007/s12975-024-01304-2.

[58] Grépin R, Guyot M, Giuliano S et al. The CXCL7/CXCR1/2 axis is a key driver in the growth of clear cell renal cell carcinoma. Cancer Res 2014;74:873–883. doi: 10.1158/0008-5472.CAN-13-1267.

[59] Alsabani M, Abrams ST, Cheng Z et al. Reduction of NETosis by targeting CXCR1/2 reduces thrombosis, lung injury, and mortality in experimental human and murine sepsis. Br J Anaesth 2022;128:283–293. doi: 10.1016/j.bja.2021.10.039.

[60] Oh SA, Seol SI, Davaanyam D et al. Platelet-derived HMGB1 induces NETosis, exacerbating brain damage in the photothrombotic stroke model. Mol Med 2025;31:46. doi: 10.1186/s10020-025-01107-7.

[61] Molinaro R, Yu M, Sausen G et al. Targeted delivery of protein arginine deiminase-4 inhibitors to limit arterial intimal NETosis and preserve endothelial integrity. Cardiovasc Res 2021;117:2652–2663. doi: 10.1093/cvr/cvab074.

[62] Wang CP, Ko GR, Lee YY et al. Polymeric DNase-I nanozymes targeting neutrophil extracellular traps for the treatment of bowel inflammation. Nano Converg 2024;11:6. doi: 10.1186/s40580-024-00414-9.

[63] Li W, Chi D, Ju S, Zhao X et al. Platelet factor 4 promotes deep venous thrombosis by regulating the formation of neutrophil extracellular traps. Thromb Res 2024;237:52–63. doi: 10.1016/j.thromres.2024.03.005.

[64] Okeke EB, Louttit C, Fry C et al. Inhibition of neutrophil elastase prevents neutrophil extracellular trap formation and rescues mice from endotoxic shock. Biomaterials 2020;238:119836. doi: 10.1016/j.biomaterials.2020.119836.

